# The aging lung mucosa: A proteomics study

**DOI:** 10.1101/2021.09.14.460375

**Authors:** Andreu Garcia-Vilanova, Angélica M. Olmo-Fontánez, Juan I. Moliva, Anna Allué-Guardia, Harjinder Singh, Robert E. Merrit, Diego M. Caceres, Jay Peters, Yufeng Wang, Larry S. Schlesinger, Joanne Turner, Susan T. Weintraub, Jordi B. Torrelles

## Abstract

The elderly population is at increased risk of acute and chronic respiratory infections and other pulmonary diseases, and it is estimated that this population will double in the next 30 years. Biochemical changes in the lung alveolar mucosa and lung cells alter local immune response as we age, creating opportunities for invading pathogens to establish successful infections. Indeed, the lungs of the elderly are a pro-inflammatory, pro-oxidative, dysregulated environment but this environment has remained understudied. We performed a comprehensive, quantitative proteomic profile of the lung mucosa in the elderly, developing insight into the molecular fingerprints, pathways, and regulatory networks that characterize the lung in old age. We identified neutrophils in the lungs of elderly individuals as possible contributors to dysregulated lung tissue environment. This study establishes a baseline for future investigations to develop strategies to mitigate susceptibility to respiratory infections in the elderly.

## INTRODUCTION

It is estimated that by 2050, the worldwide population of individuals who are 65 years and older will have nearly doubled to 22% of the population [1]. The elderly population is at an increased risk for developing respiratory infections, autoimmune and neoplastic diseases [2]. Significant to this study, lower respiratory tract infections and other lung pathologies are some of the top causes of death globally, independent of income group [3].

The process of aging is associated with innate and adaptive immune dysfunction and dysregulation, increased basal inflammation, and cellular metabolic dysregulation, a process commonly referred to as “inflammaging,” characterized by cellular senescence and immunosenescence [4–6]. This process inevitably leads to loss of tissue homeostasis, cellular component dysfunction, and altered protein processing and transport, and contributes to reduced capacity to mount an effective immune response, leading to increased susceptibility to respiratory infections in the elderly [7–9].

The lung alveolar mucosa, which is composed primarily of a surfactant lipid layer and an aqueous hypophase rich in soluble proteins, defined as Alveolar Lining Fluid (ALF), is constantly produced and recycled in the lung by the Type II alveolar epithelial cells (AT-IIs) [10, 11]. We hypothesized that “inflammaging” negatively alters the phenotype and function of AT-IIs and other lung structural and alveolar-resident and -circulating compartment cells. Negative alterations can occur at the cellular level in the form of aging-directed immune cell dysfunction and tissue level through oxidative changes to the alveolar environment. These changes inevitably lead to the composition and function of the alveolar mucosa and the local innate immune responses. This is of special relevance, as this is the site of the first contact between external pathogen and host cells.

Our previous studies have demonstrated that the elderly lung constitutes a highly pro-oxidative and pro-inflammatory environment, with altered levels of complement and surfactant proteins, as well as the decreased binding capacity of surfactant protein A (SP-A) and D (SP-D) [12]. As a consequence, exposure of the intracellular pathogen *Mycobacterium tuberculosis* (*M.tb*) to elderly ALF results in subsequent increased *M.tb* intracellular growth *in vitro* (in macrophages and ATs) and *in vivo* (mouse model) upon infection [13, 14]. Understanding the changes that shape this susceptibility phenotype in the elderly population is imperative to develop preventative care strategies and treatments to improve care for the elderly.

In-depth studies of “inflammaging” and the biochemical status of ALF in the elderly human population are almost non-existent, especially in healthy individuals, due to either the difficulty in obtaining viable samples and the cost of acquisition and analysis of such samples [15, 16]. Nonetheless, understanding the biochemical composition of ALF in elderly individuals should provide valuable insight into the changes that occur during aging in the lung, which predisposed the elderly to respiratory infections and pathologies.

In this study, we compared the proteomic profiles of ALF from healthy elderly humans (62 and older) to ALF from healthy younger individuals (48 and under). We identified individual molecules, networks, and pathway regulators that are significantly altered in the ALF from elderly individuals. We discuss how these differences may be responsible for the susceptibility to respiratory infections observed in the elderly.

## RESULTS

### Comparative mass spectrometry-based proteomic analysis of human E-ALF *vs*. A-ALF

We performed Data-independent acquisition mass spectrometry (DIA-MS) to obtain identifications and relative quantifications of proteins in human ALF from six adults (48 and under) and ten elderly (62 and older) healthy donors. A total of 2,239 proteins and 14,844 peptides were identified (1,771 proteins with a minimum of two peptides quantified). We identified Differentially Expressed Proteins (DEPs) from the elderly group that we deemed biologically relevant for this exploratory study, defining p < 0.05 based on the Benjamini-Hochberg multiple corrections control of FDR set at 20% [17], and log_2_-transformed fold change ≤ 180 cl:1300.5 and ≥ 0.5, which yielded a total of 457 DEPs of interest (**Fig. 1**).

**Figure 1.**
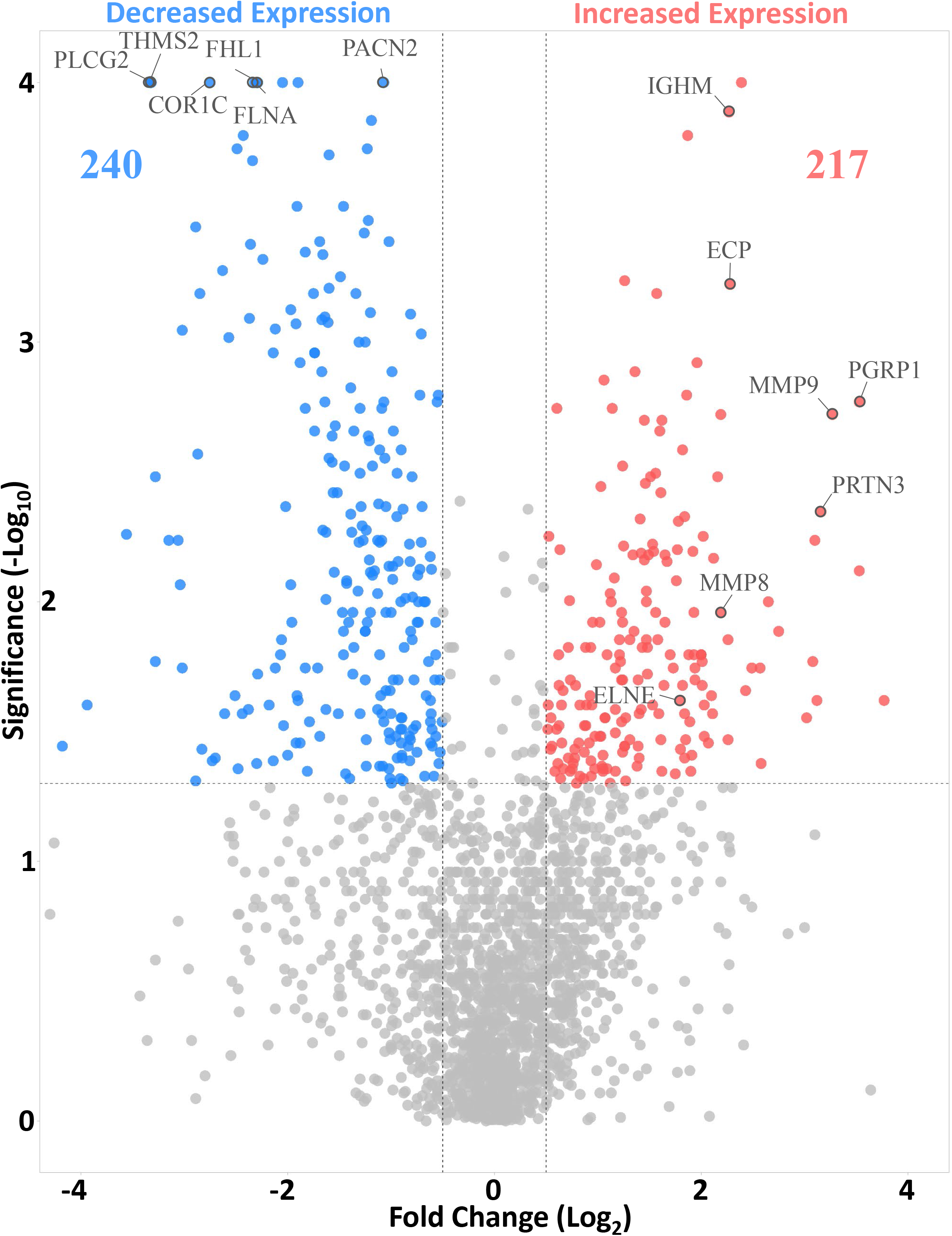
Volcano plot of the distribution of Differentially Expressed Proteins (DEPs). The x-axis shows log_2_-transformed protein fold changes for human E-ALF *vs*. A-ALF, the Y-axis shows −log_10_-transformed *p*-values. A total of 457 DEPs were identified, 217 being higher (Red) and 240 being lower (Blue) in E-ALF.

A total of 217 DEPs were found to be significantly higher in elderly ALF (E-ALF), while 240 DEPs to be significantly lower when compared to adult ALF (A-ALF). Of the 217 DEPs significantly higher in E-ALF (**Table 1,** Log_2_ fold changes), 156 were at *p*<0.05; 46 at *p*<0.005; 9 at *p*<0.0005 (e.g., ASSY, CALU, CAN2, CAPS2, CPNS1, NASP, PAPS1, PGRP1, and SPB10), and six at *p*<0.0001 (e.g., ECP, IGHM, MMP9, NCHL1, RSH4A, and S100P). Of the 240 significantly lower in E-ALF (**Table 2,** log_2_ fold changes), 133 were at *p*<0.05; 76 at *p*<0.005; 23 at *p*<0.0005 (e.g., ARC1B, ARP2, ARP3, ARPC5, CSRP1, DIAP1, EMAL4, IFTI3, IST1, KPCD, MK14, MOES, NCF4, PANC2, PLCG2, PLEK, SARG, SNX5, SYVC, UPP1, VAT1, VATE1, and URP2), and eight at *p*<0.0001 (e.g., BAX, CLIC4, COR1C, FHL1, FLNA, FLNC, PTN6, and THMS2). In E-ALF samples, 28.1% of highly significant proteins were higher, while 44.6% were lower when compared to A-ALF samples, suggesting a skew towards reduced production. Of the total proteins identified, approximately 30% were annotated as being intracellular.

**Table 1.**
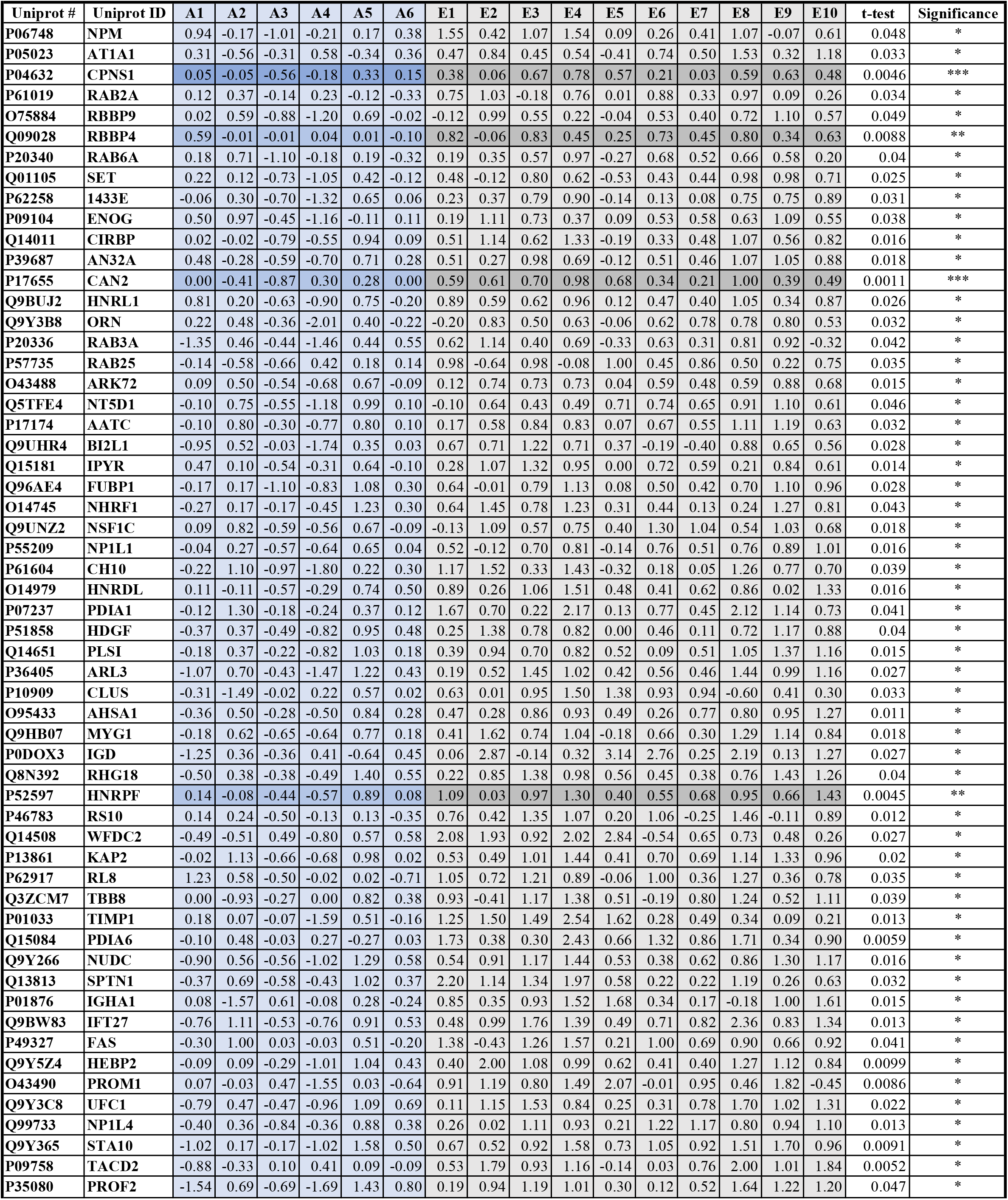
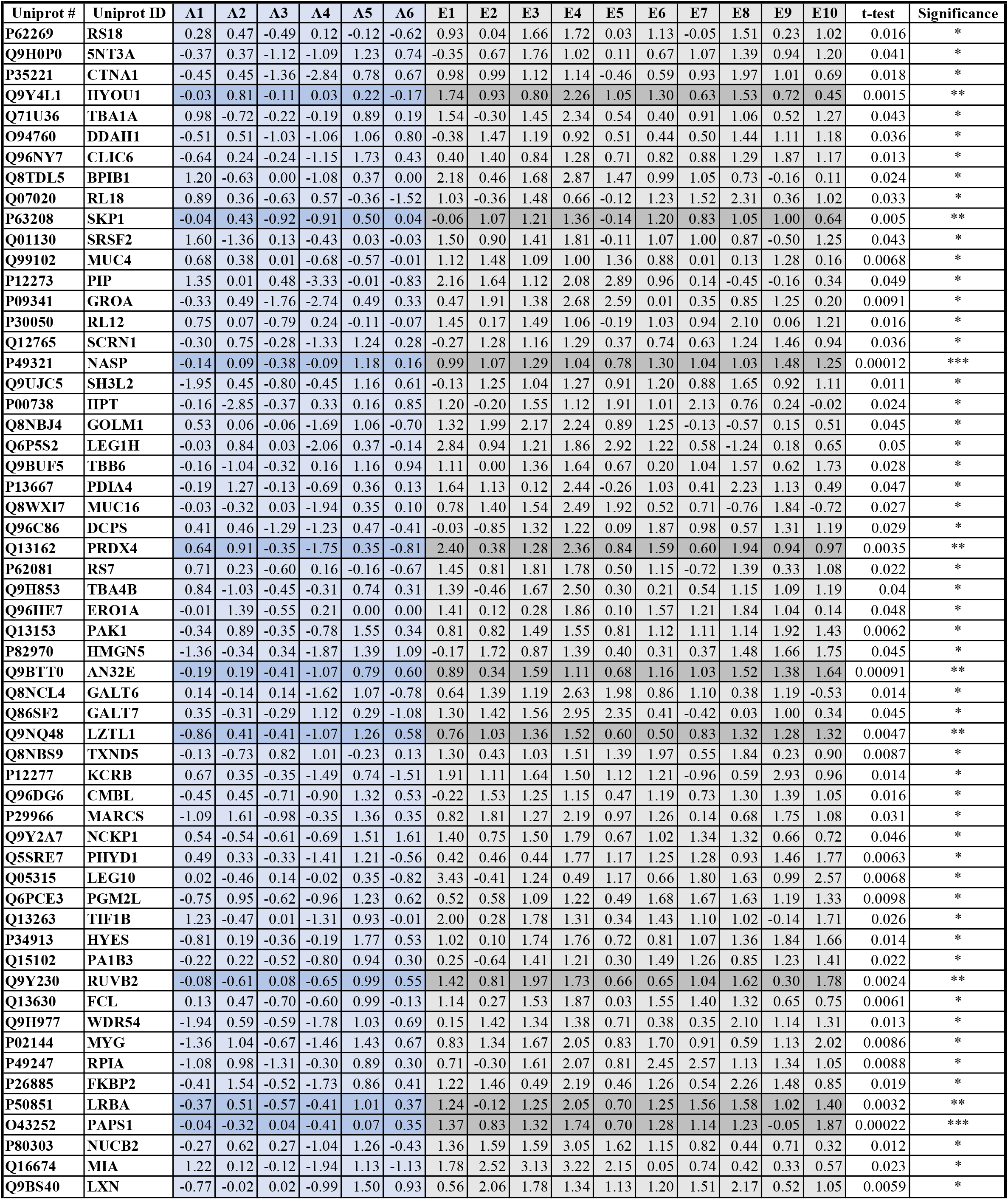
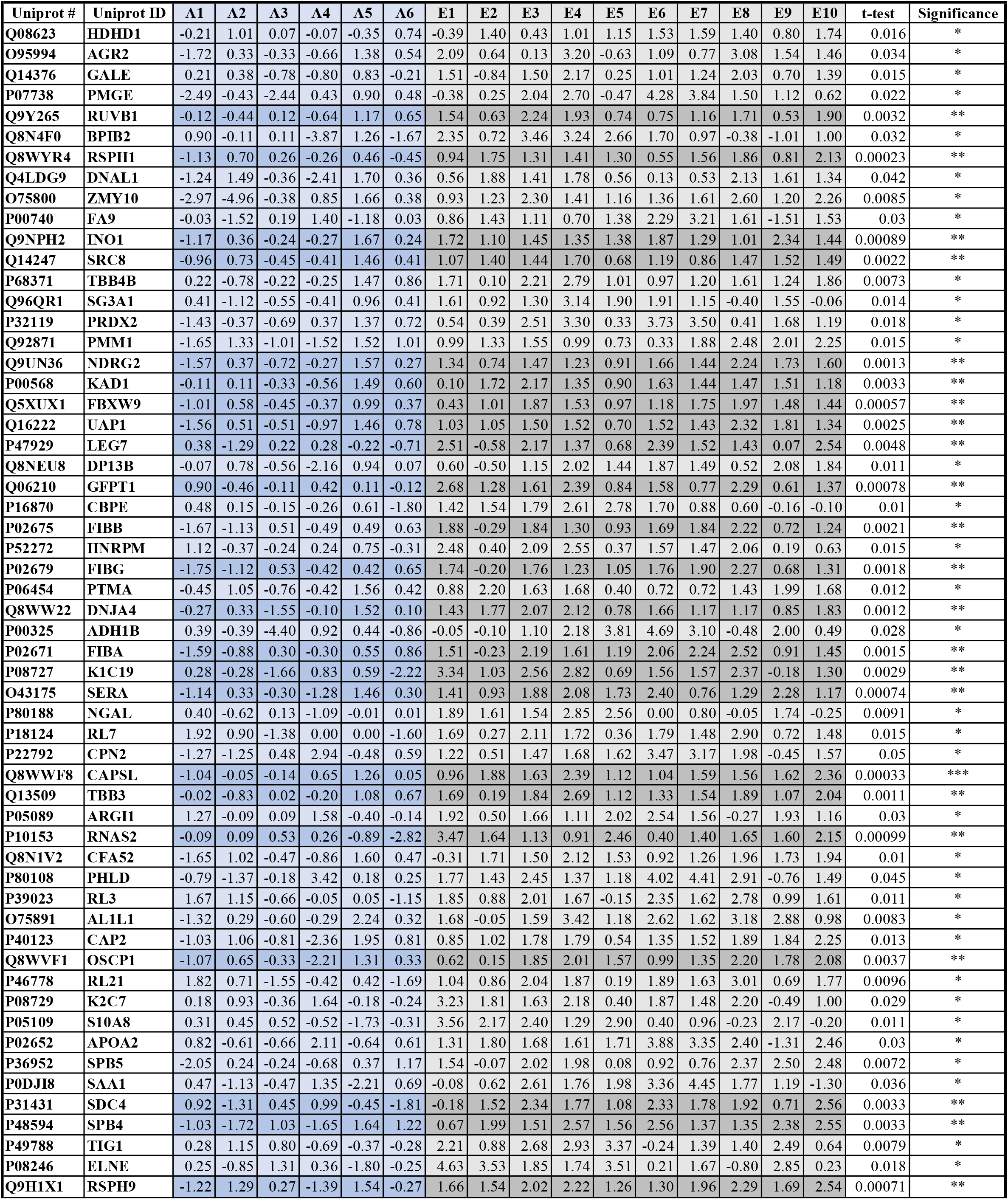
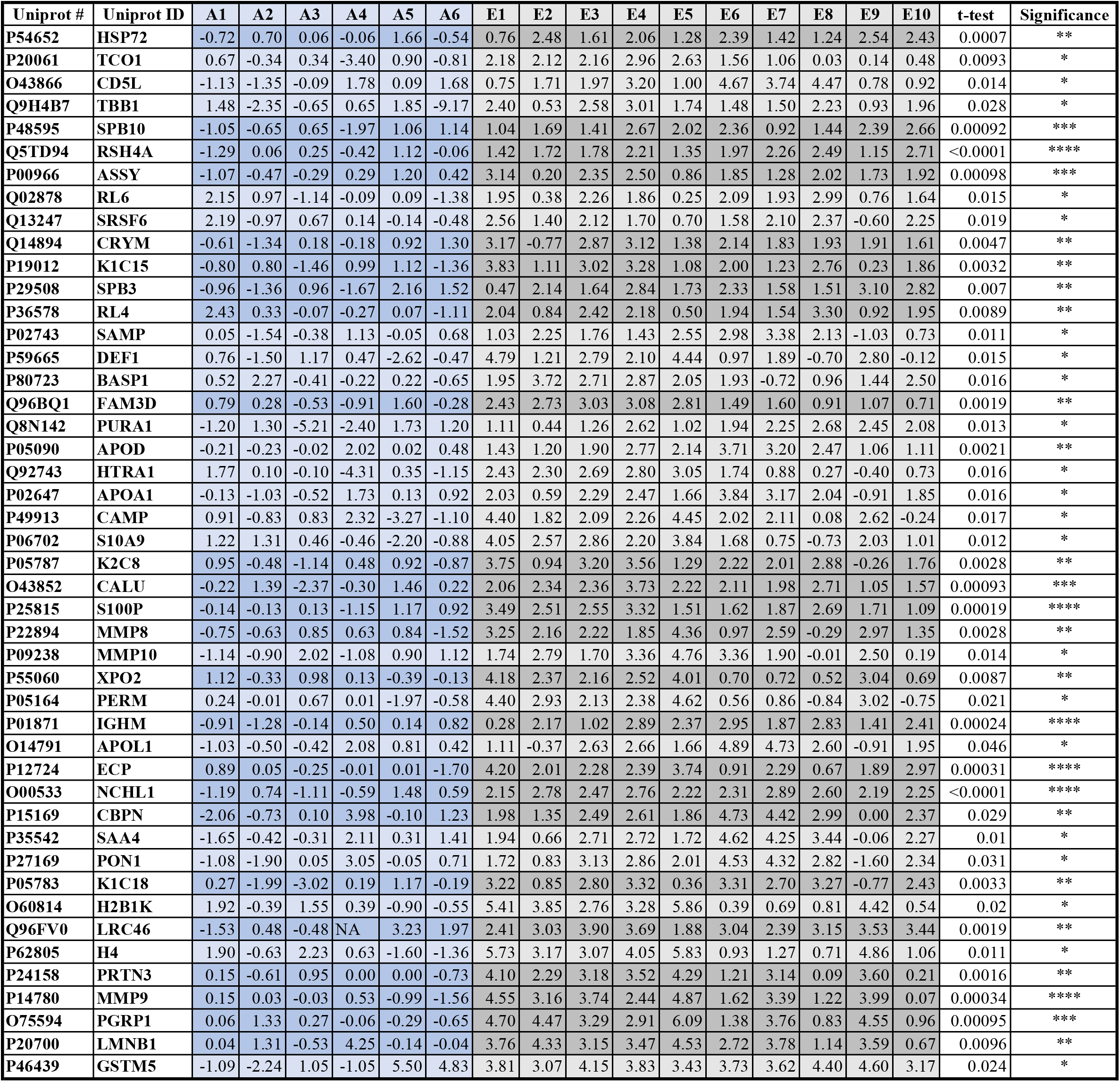
Detailed log_2_-transformed Fold Changes for higher DEPs in the Elderly. A summary table of proteins significantly higher in E-ALF vs. A-ALF. \**p*<0.05; \*\**p*<0.005; \*\*\**p*<0.0005; \*\*\*\**p*<0.0001, ANOVA with Benjamini Hochberg control of FDR at 20%. Highlighted darker lines in for lower p-value group.

**Table 2.**
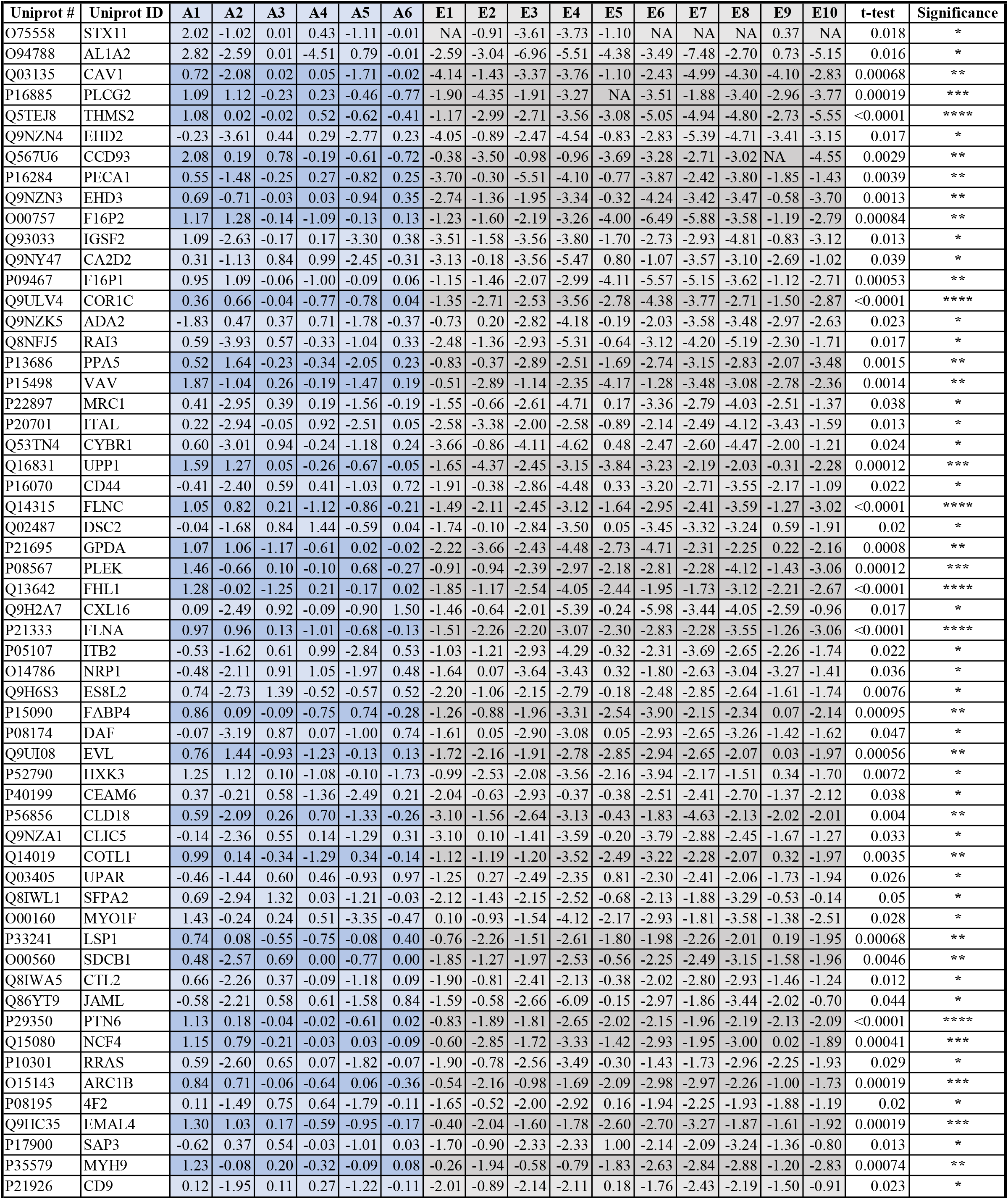
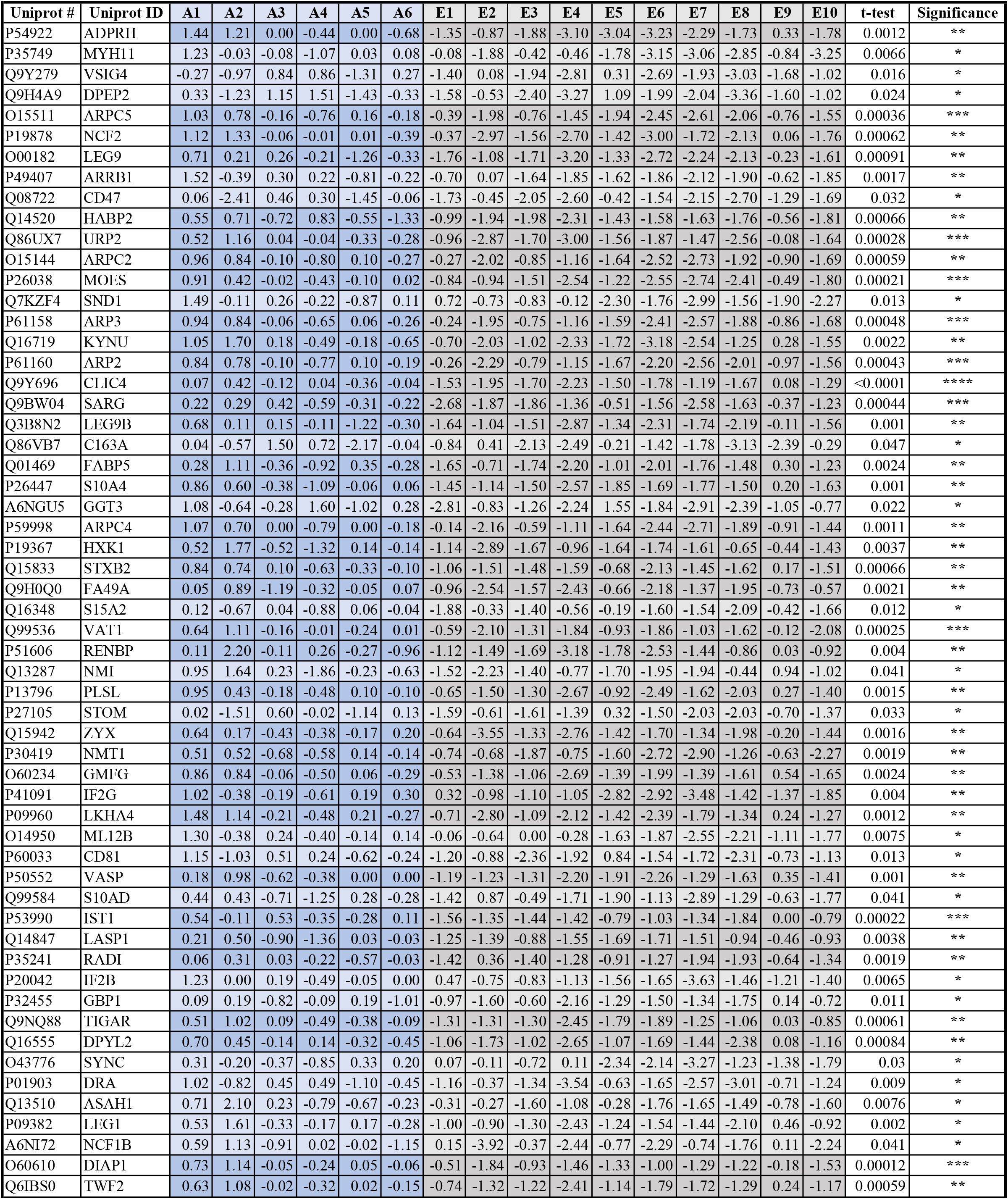
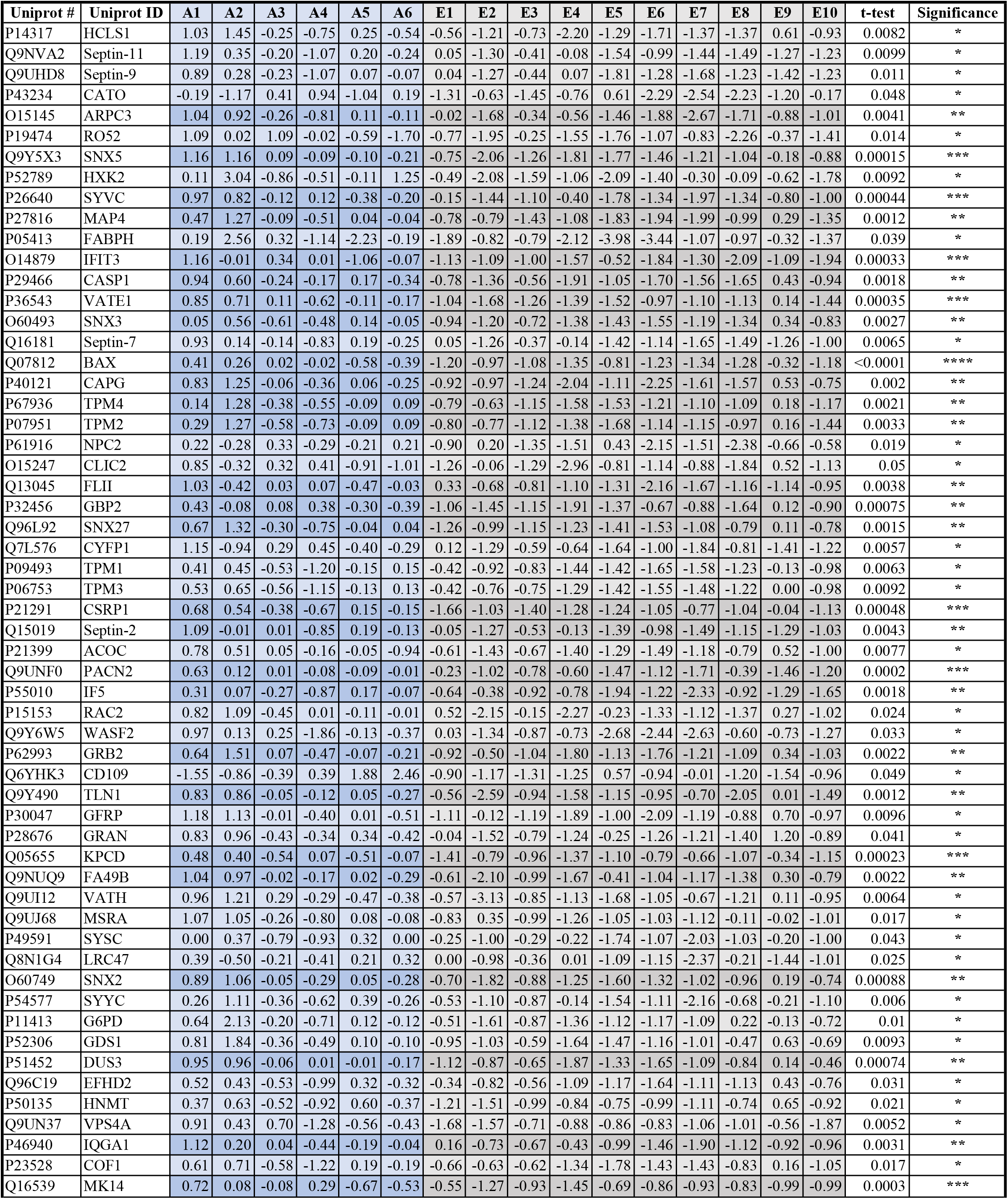
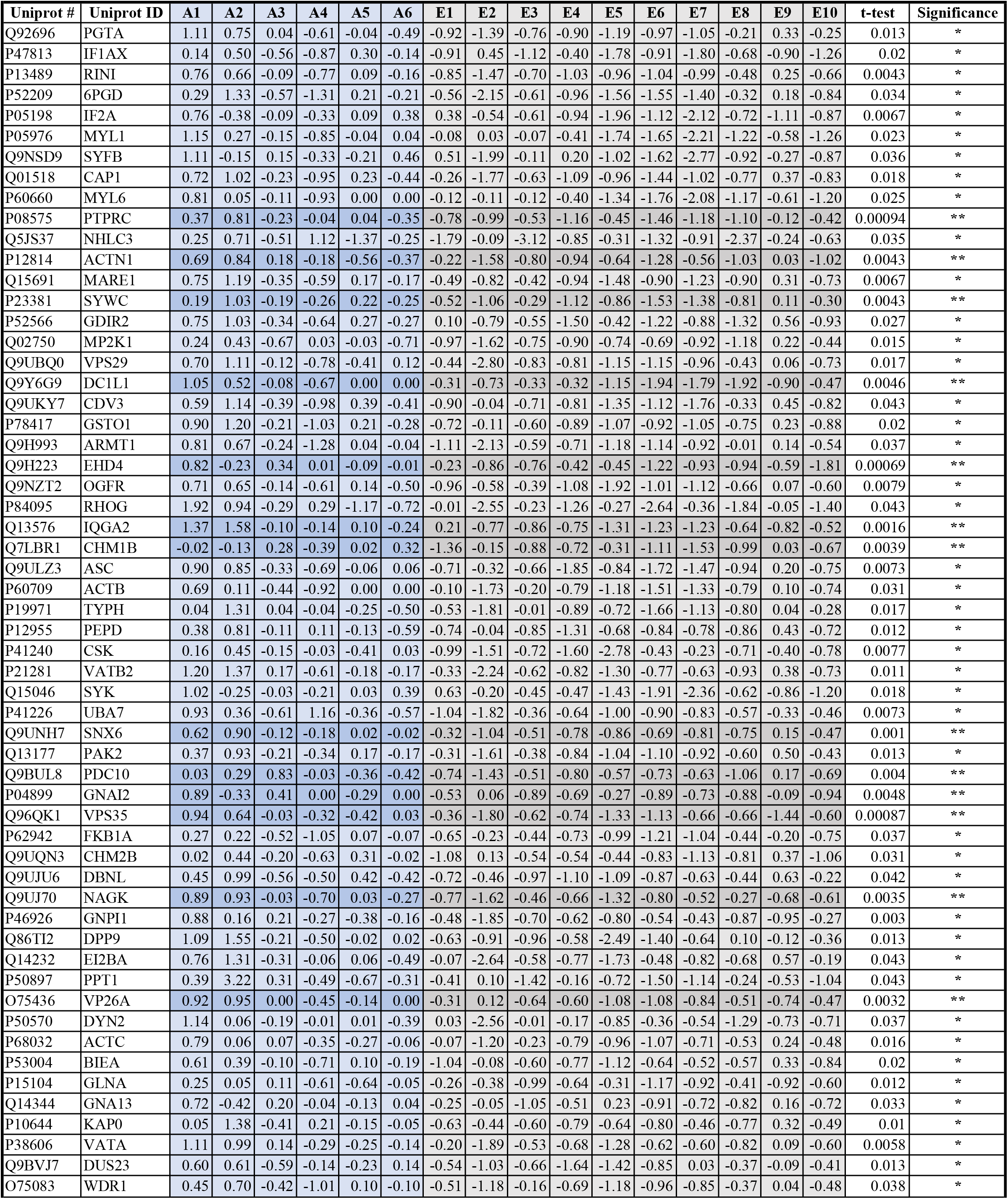
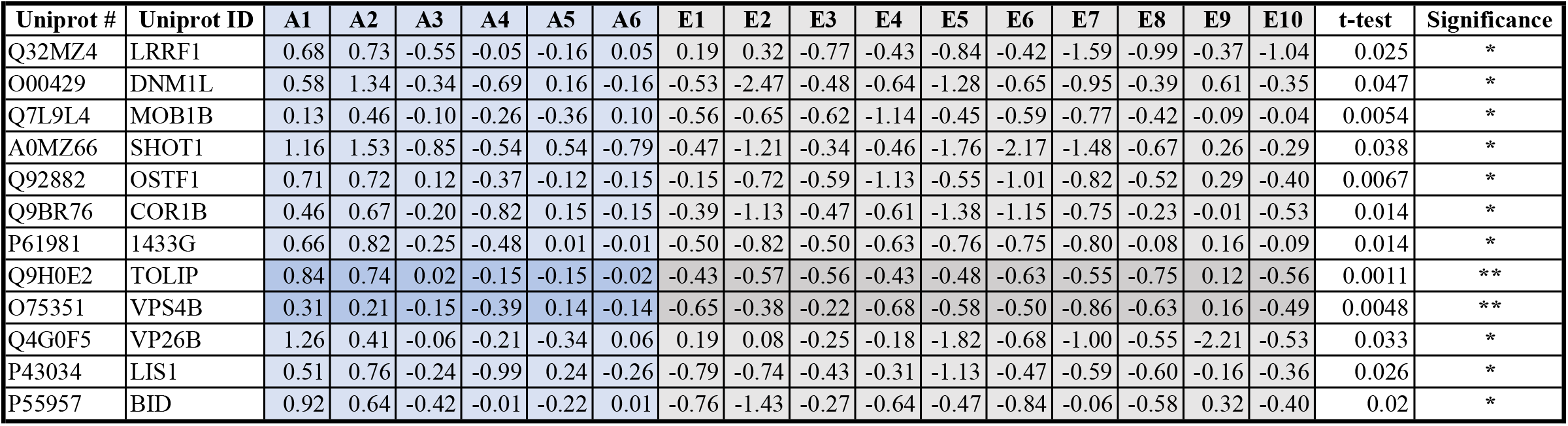
Detailed log_2_-transformed Fold Changes for lower DEPs in the Elderly. A summary table of proteins significantly lower in E-ALF vs. A-ALF. \**p*<0.05; \*\**p*<0.005; \*\*\**p*<0.0005; \*\*\*\**p*<0.0001, ANOVA with Benjamini Hochberg control of FDR at 20%. Highlighted darker lines in for lower p-value group.

We performed a hierarchical clustering analysis to determine how samples were distributed (**Fig. 2**). A-ALF samples had higher variability among donors than the E-ALF samples, which clustered tightly. ALF A5 clustered with the E-ALF group and was, together with E9, the more different sample in the E-ALF cluster. This result was somehow expected given the sample size and the intrinsic variation among healthy humans. Further analysis of the human ALF donor demographics showed no correlation between sample groupings with regard to race or sex (data not shown), pointing to age as the most likely factor to explain the variability across E-ALF *vs*. A-ALF.

**Figure 2.**
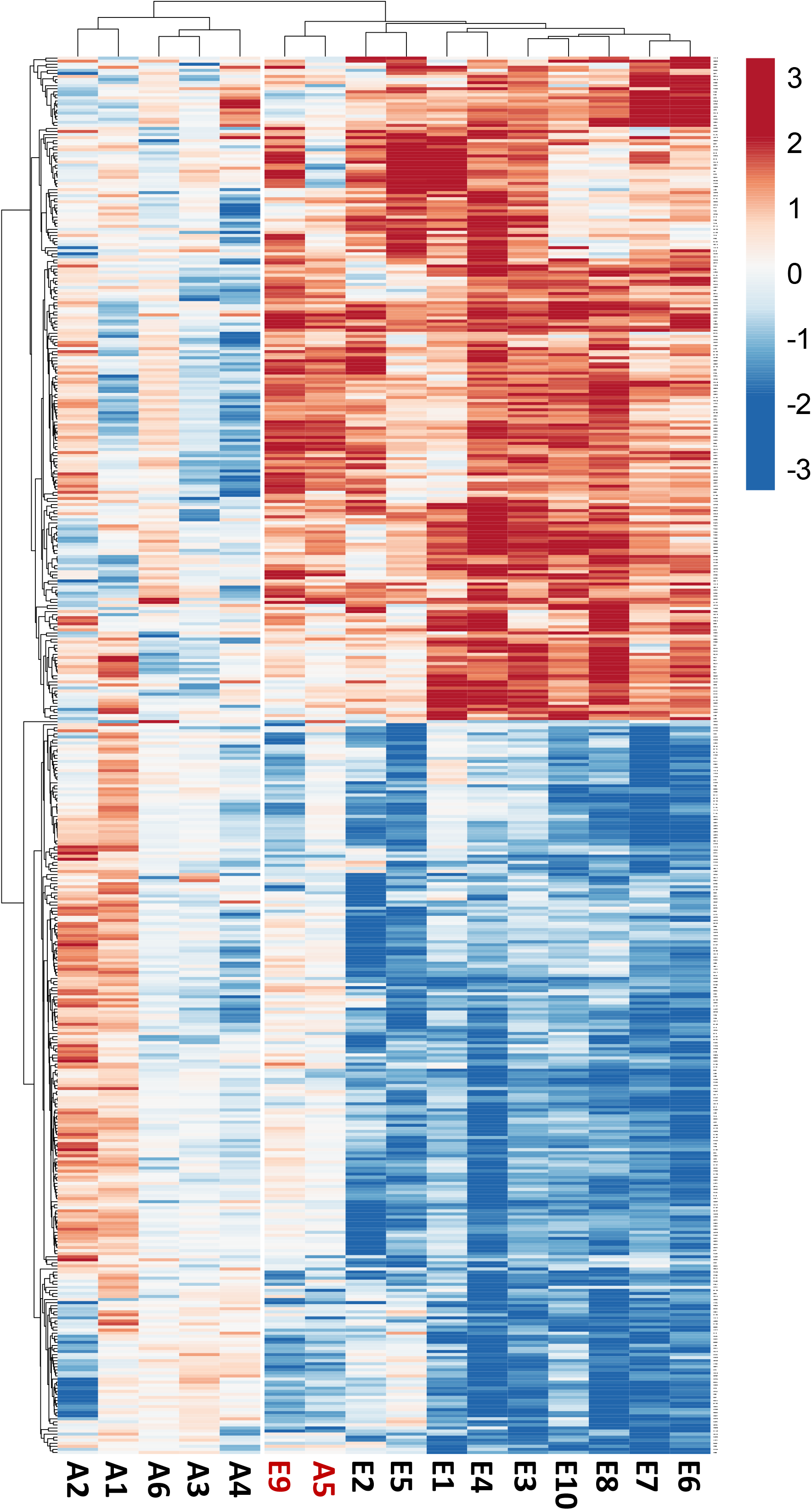
Hierarchical clustering of DEPs. Heatmap showing sample number on the X-axis, Uniprot Accession numbers on the Y-axis, and log_2_-transformed fold changes (Z, Blue [lower] to Red [higher]). Clustering analysis reveals higher variability among the A-ALF samples (adults), with E-ALF samples (Elders) strongly clustering with each other. Samples A5 and E9 were identified as the most different from their respective groups.

### Gene Ontology (GO) enrichment analysis of human E-ALF *vs*. A-ALF

GO annotations and classifications were determined for the 217 and 240 identified proteins that were found to be significantly higher or lower in the E-ALF *vs*. A-ALF (**Fig 3**). The enriched GO terms with over 250 proteins assigned under the biological process category were cellular processes, biological regulation, and response to stimulus. The GO annotations with over 250 proteins assigned within the cellular component category were cytoplasm, intracellular organelle, extracellular region, organelle part, and membrane. Lastly, categories with over 150 proteins were classified within the binding and catalytic activity annotations for the molecular function category. A summary of the GO annotations for all detected proteins regardless of their significance between E-ALFs and A-ALFs is provided in **Supplemental Fig S1**.

**Figure 3.**
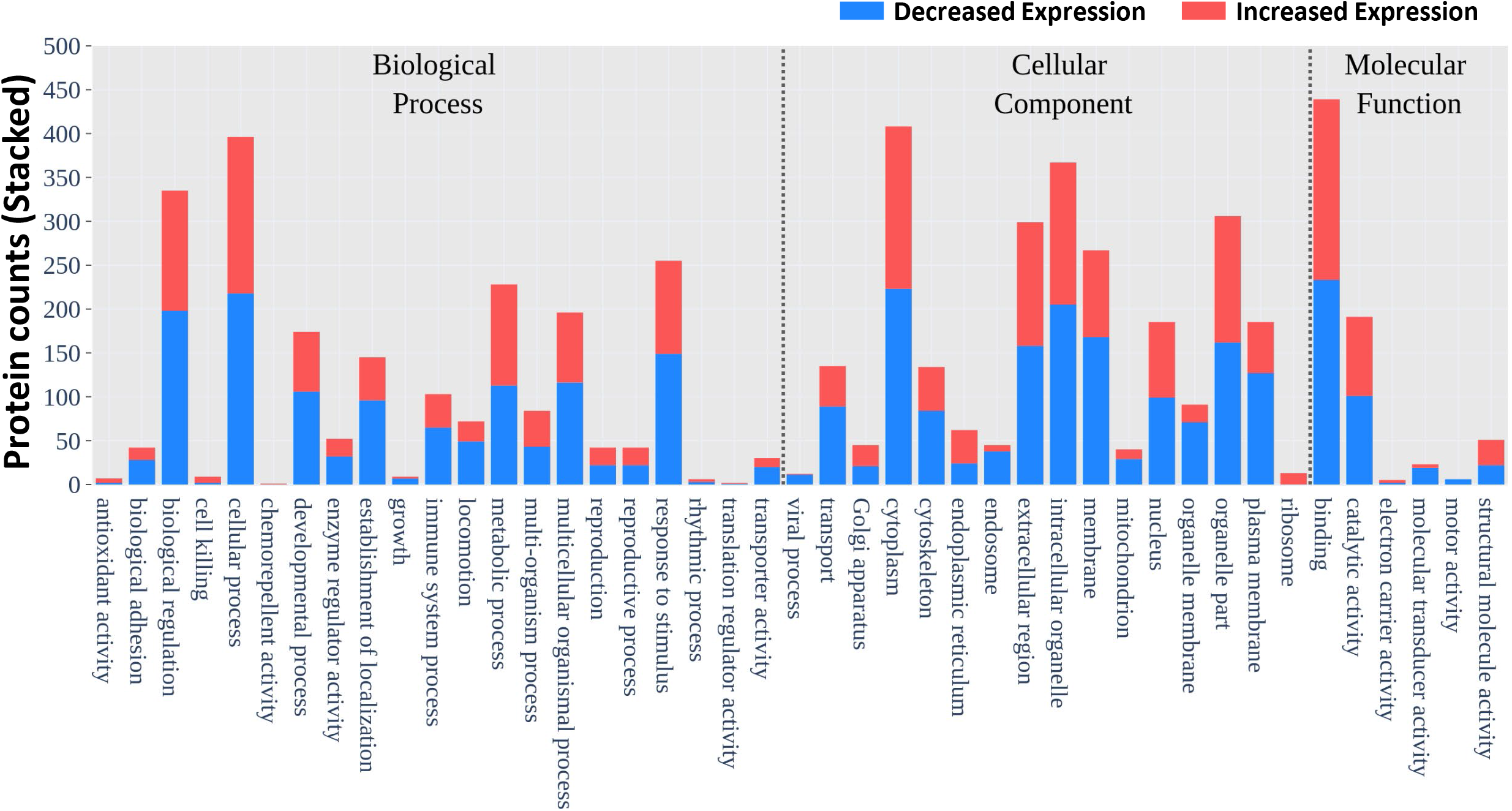
Gene Ontology annotations, distribution of DEPs. Stacked bar chart of the total number of DEPs (Blue [Lower] and Red [Higher]) belonging to the different GO classifications (Biological Process, Cellular Component, and Molecular Function).

### Top identified canonical pathways enriched in human E-ALF *vs*. A-ALF

We further used Ingenuity Pathway Analysis (IPA, Qiagen) to identify the top canonical pathways overrepresented by the identified DEPs, the top 40 significantly overrepresented pathways in E-ALFs compared to A-ALF are shown (**Fig. 4)**. As seen in the figure, the top 40 pathways (selected by the highest p-values of overlap) had a high overlap, as defined by the right-tailed Fisher’s exact test (−Log p-values > 4.02), color-coded in the figure. The pathways with the highest −log p-values were ‘Actin Cytoskeleton Signaling,’ ‘Remodeling of Epithelial Adherens Junction,’ and ‘Integrin Signaling.’ Values for the ratio of pathway proteins (DEPs in dataset belonging to pathway / Proteins in the pathway for reference dataset) is provided as the dot size, with the highest ratios found for the ‘UDP-N-acetyl-D-galactosamine Biosynthesis II,’ ‘Remodeling of Epithelial Adherens Junction,’ and ‘tRNA charging.’ Finally, the activation z-score, a prediction of the overall activation state of a pathway based on the status of the detected proteins, is shown by the bubble position on the x-axis, with the biggest changes being ‘Signaling by Rho Family GTPases’ and ‘RHOA Signaling,’ for the inhibited pathways, and ‘RHOGDI Signaling’ as the top activated pathway. A table of the top 100 pathways identified by IPA is also provided, selected by their p-values (**Table 3**).

**Figure 4.**
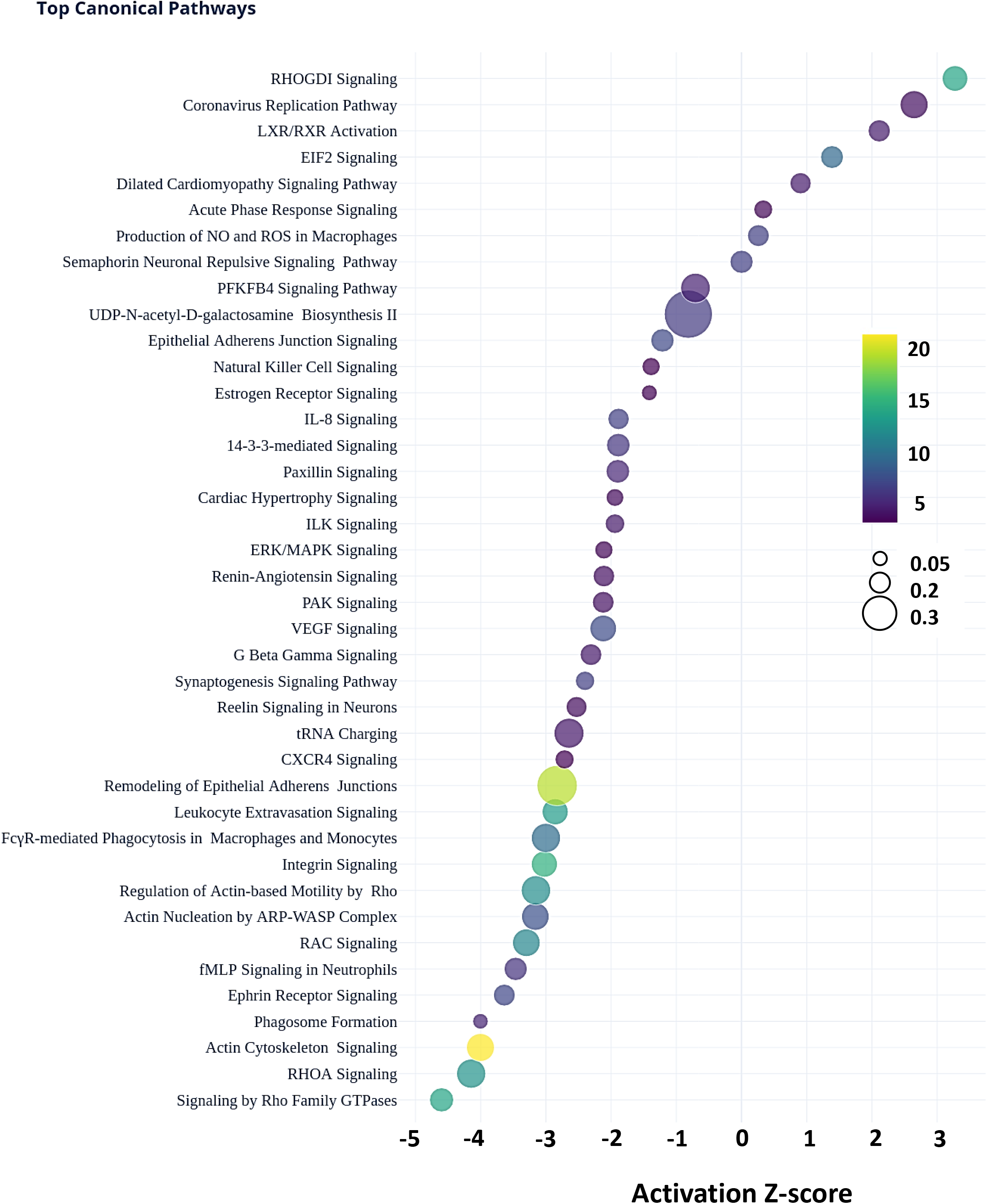
Top canonical pathways. A bubble plot summarizing the top 40 canonical pathways of the DEPs. Pathway activation Z-scores, defined by bubble position on the X-axis (−4.146 – 3.273), Pathway name on the Y-axis, −log-transformed p-values (Color scale, blue-to-yellow, 4.02 – 22.8), and Ratio (DEPs in dataset belonging to pathway / Proteins in the pathway for reference dataset, bubble size, small-to-big, 0.0479 – 0.462).

**Table 3.**
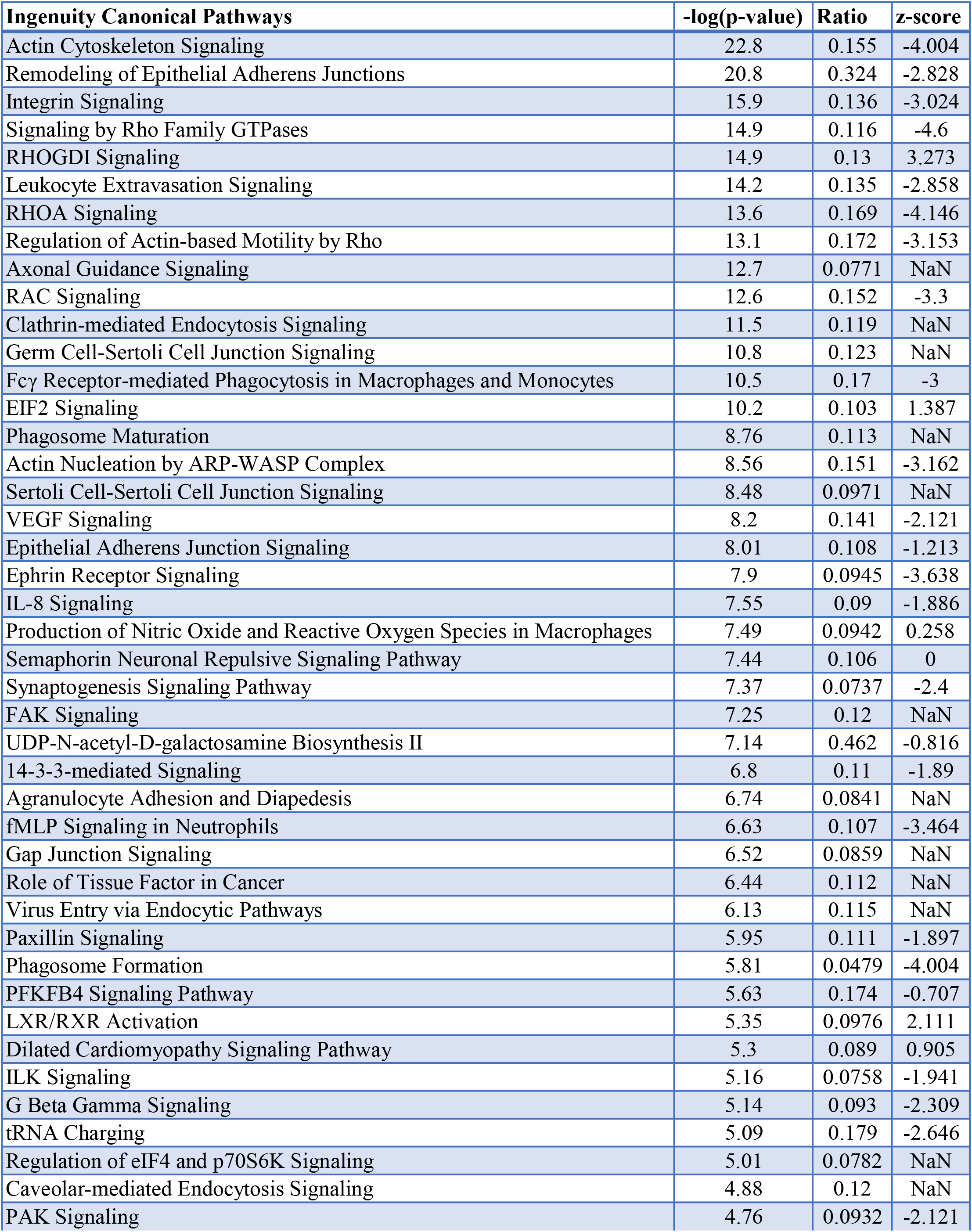
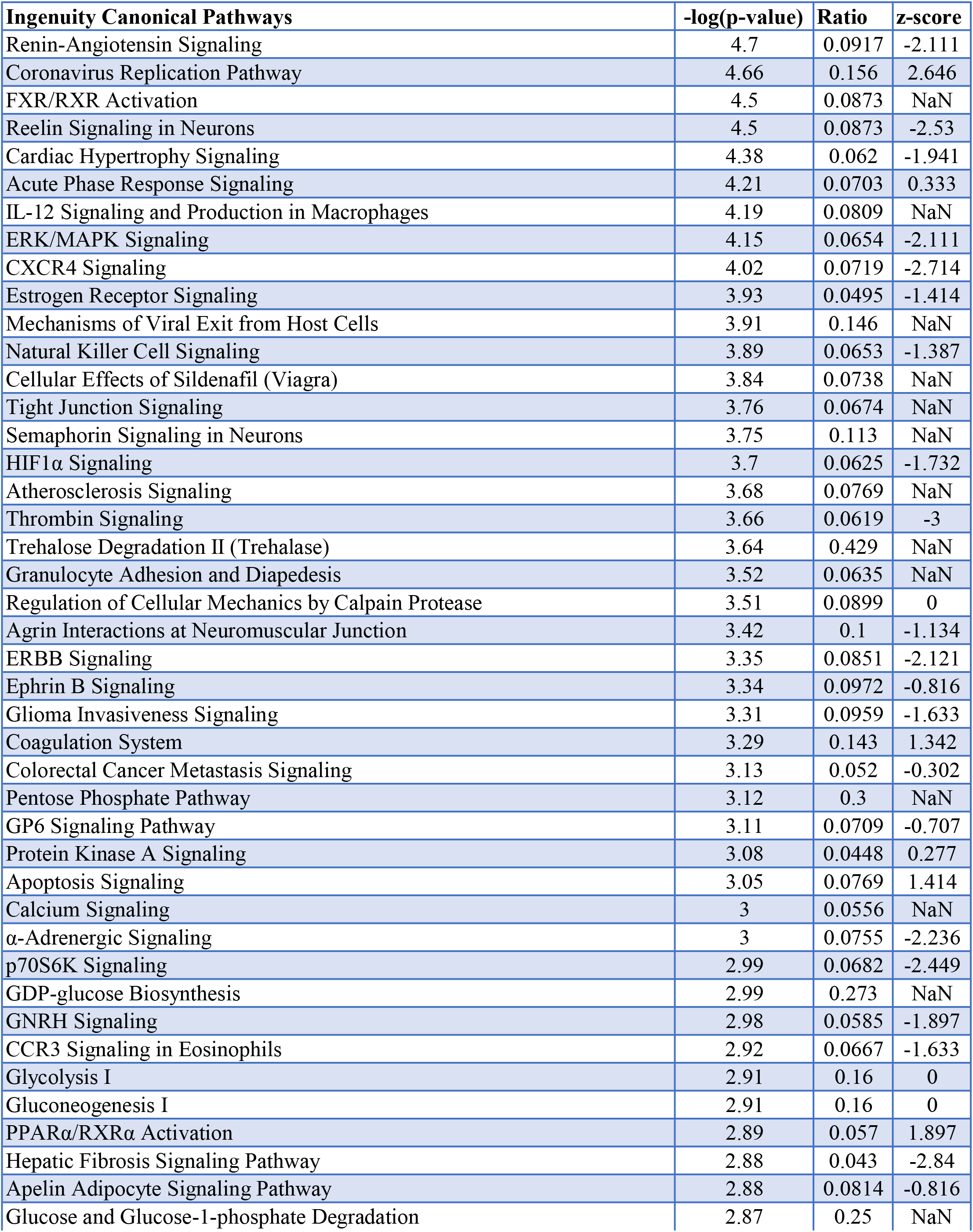
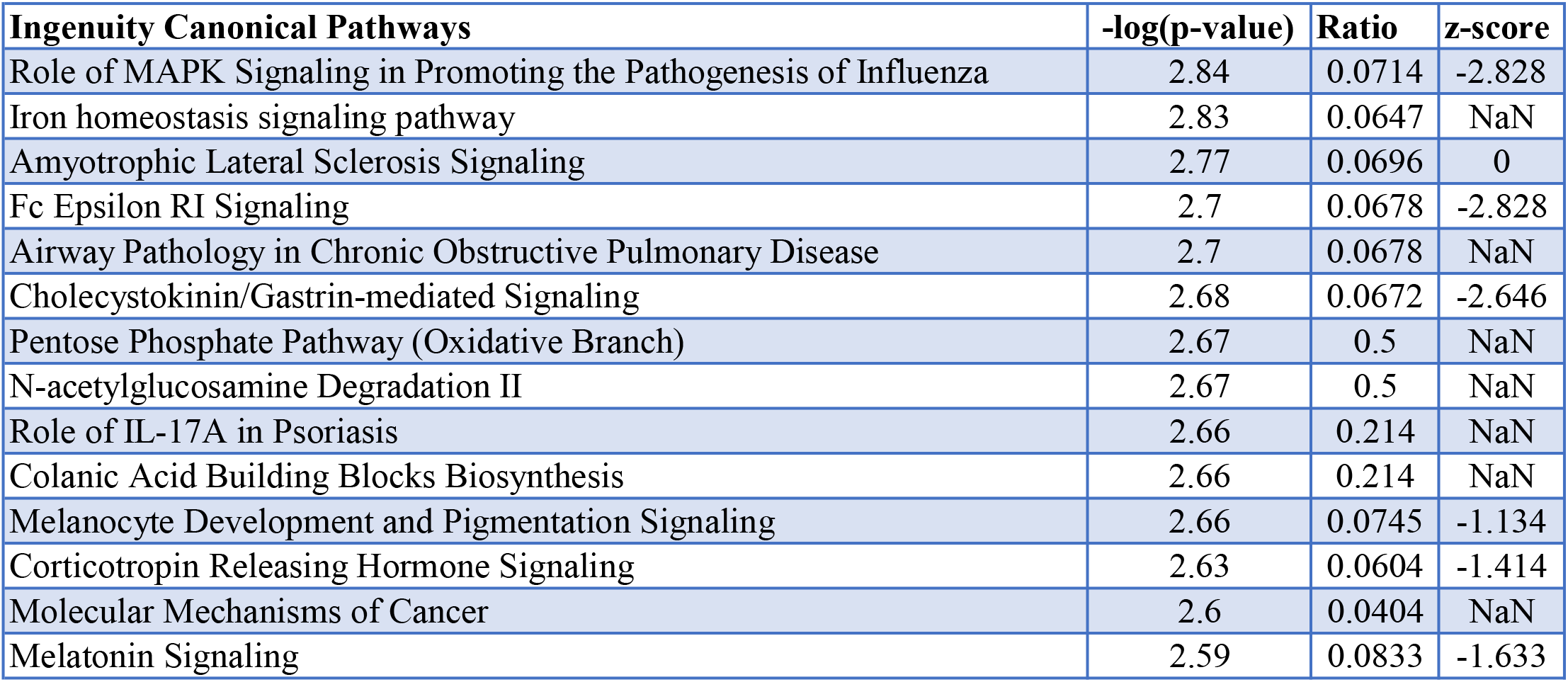
Summary of the Top 100 Canonical Pathways. A summary table of significantly enriched Canonical Pathways identified by IPA analysis, showing −log(p-value) of the protein overlap, a ratio of DEPs in dataset belonging to pathway / Proteins in the pathway for reference dataset, and an activation z-score, an estimation of the overall activation of the pathway based on the status of the molecules identified in E-ALF.

## DISCUSSION

The lung is one of the major organs in constant direct contact with the external environment, and as a result, has evolved to be unique in terms of cell metabolism, tissue organization, and immune responses [18, 19]. The lung mucosa is specialized with three main functions: maintenance of alveolar structure by preventing alveolar collapse (by surfactant lipids), facilitation of gas exchange between the external air and the alveolar epithelial cells, and serves as the first line of defense against external insults, which is the primary function of the ALF hypophase portion of the lung mucosa [11, 20, 21]. Our laboratory has previously demonstrated that ALF can shape responses to microbial infections [22–27], and that dysfunctional soluble proteins encountered in the elderly ALF were associated with worse infection outcomes, both *in vitro* and *in vivo* [12–14]. We investigated the proteomic make-up of E-ALF *vs*. A-ALF to better understand age-related changes that occur in the soluble proteins of the lung and to gain an insight into changes that may occur at the cellular level, capturing intracellular proteins in the ALF compartment [28]. Our results characterize proteins that are significantly altered in their presence in the elderly ALF. Of relevance to this discussion, approximately 30% of proteins identified in the alveolar lining fluid were considered to be intracellular. The presence of intracellular proteins in the ALF could be explained by several factors, including but not limited to cell contents released to the ALF after cell death by necrosis [29], increased cell turnover [30], proteins contained in the lumen of secreted vesicles [31], or secreted in lamellar bodies by ATs [32]. It is important to note that the mechanisms by which some intracellular proteins are found in the extracellular space are not entirely understood, and extra precautions need to be taken before raising any conclusions. In this study, the findings discussed below related to intracellular proteins identified are purely speculative but of importance nonetheless, as those were significant changes detected in the E-ALF.

A protein class detected at higher levels in E-ALF was the matrix metalloproteinase (MMP) family (**Tables 1 and 3**) and their regulatory counterparts, the tissue inhibitors of matrix metalloproteinases (TIMPs). MMPs are a large family of Zn^2+^- and Ca^2+^-dependent endopeptidases produced by several cell types [33, 34]. Increased levels of MMPs strongly correlate with chronic inflammation and tissue destruction, hallmarks of lung catabolism and diseases [35]. We detected four different MMPs, MMP-8, −9, and −10, significantly higher in the elderly (2.2, 3.41, and 2.19 Log_2_ fold change, respectively), and MMP-7, although not significant, showed a trend towards overproduction in the elderly. We also detected TIMP1 (0.87, Log_2_ Fold change), a metalloproteinase inhibitor that acts on all four detected MMPs [36]. Although not statistically significant, the major inhibitor of MMP9, α-2-macroglobulin [37], showed a trend towards overproduction. This profile of increased extracellular matrix remodeling, among others observed, suggests a chronic immunoinflammatory status of the lung of elderly individuals.

The significantly higher production of proteins of neutrophilic granule origin, such as mainly azurophil granule proteins like proteinase 3 (PRTN3), myeloperoxidase (PERM), elastase (ELNE), and defensin-1α (DEF1), and specific granule proteins like cathelicidin (CAMP) and gelatinase-associated lipocalin (NGAL), support the role of a neutrophilic contribution to the overall pro-inflammatory and pro-oxidation status of the lung mucosa in the elderly. It remains unclear whether neutrophils in the lung of elderly individuals undergo senescence in the alveoli in the same manner as other cell types do.

However, it has been noted that neutrophils isolated from the elderly have altered function and increase damage to tissues [38], and the lungs of the elderly harbors increased numbers of neutrophils [39, 40]. More importantly, the neutrophil phenotype in elderly individuals could be one of the initiators of cellular senescence in the lungs by inducing neutrophilic damage to telomeres of resident and compartment cells [41]. To further examine this, we looked for regulators that could drive the neutrophilic phenotype in the lungs of elderly individuals. Our Causal Network Analysis in IPA looking at identified proteins of neutrophilic origin provided insight into a regulatory network orchestrated by the serum response factor (SRF), predicted to be underproduced in the lungs of the elderly based on our downstream data. SRF is a transcriptional factor associated with dysregulation of neutrophils and has been suggested to be altered in the elderly [42, 43]. We present a theoretical regulatory network for the neutrophil phenotype observed in our analysis, consisting of the main predicted regulator, SRF, together with its two cofactors, Myocardin Related Transcription Factors A and B (MRTFA & MRTFB), and the predicted indirect regulator Rho GTPase activating protein 4 (ARHGAP4) (**Fig. 5)**.

**Figure 5.**
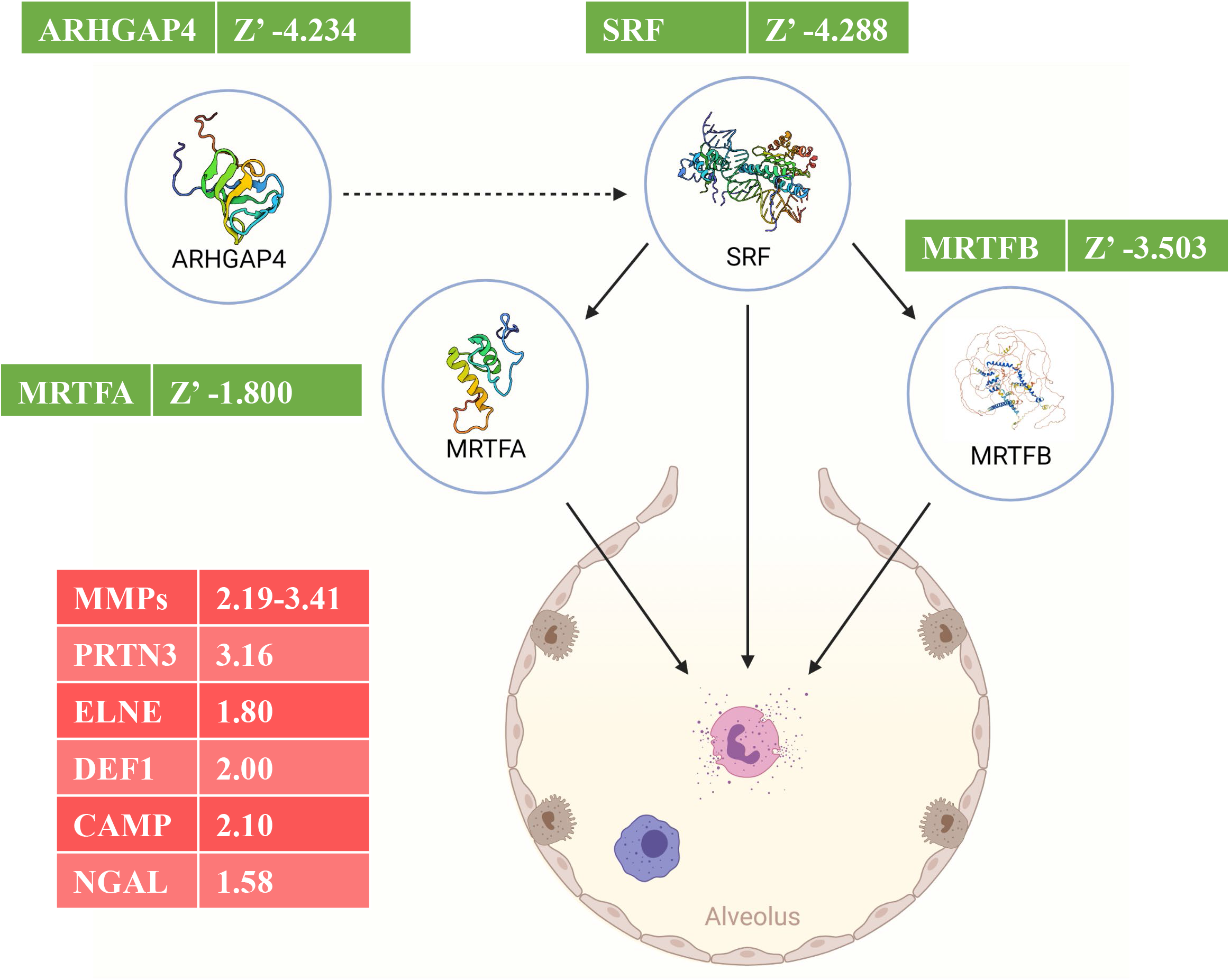
Predicted regulatory network. A regulatory network showing the different proteins that could be playing a role in downstream effects of excess neutrophil degranulation in E-ALF. Each molecule is accompanied by higher detected log_2_ fold changes (Red), or lower predicted activation Z-scores (Green). This illustration was created with BioRender (https://biorender.com/).

In our analysis, Lamin B1, a biomarker of cellular senescence, was found to be highly produced in E-ALF (**Table 1**); however, its high production is contrary to other reports showing lower production in aging [44–46]. The Peptidoglycan Recognition Protein 1 (PGRP1) also appeared significantly elevated in E-ALF. PGRP1 plays a bactericidal function in antimicrobial defense systems, acting as a pattern receptor for murein peptidoglycans (PGN) of Gram-positive bacteria [47]. Interestingly, significantly elevated levels of eosinophil cationic protein (ECP, p<0.0005), present in eosinophil granulocytes resident in lung mucosa [48], could indicate degranulation of these cells in the lungs of the elderly, driving increased inflammatory cell functions in the lung, which could result in eosinophilic disease [49]. Like PGRP1, other proteins elevated (p<0.0005) in E-ALF previously found to be involved in early recognition of external pathogens, including lung antimicrobial ciliary mobility and microvilli formation in lung epithelial cells, were IGHM (Immunoglobulin Heavy Constant Mu) and S100P (S100 Calcium Binding Protein P).

Based on the intracellular proteomic footprint found in E-ALF derived from alveolar resident and compartment cells, some of the top pathways identified were closely related to actin dysfunction in the elderly (**Fig. 4).** Of specific importance, we identified proteins that may indicate a decrease in actin polymerization of cells in the elderly, *i.e*., lower levels of ARP2/3 complex, including APR complex (APRC) subunits, 2, 3, 4, and 5; and lower actinin 1 (ACTN1) and ACTB/C [involved in polymerization of globular actin (G-actin) to form a structural filament (F-actin)] [50], and DIAP1; driving a decrease in actin stabilization and depolymerization, evidenced by lower levels of Cofilin (COF1). ARP2/3 complex regulates the actin cytoskeleton, where two of its subunits (ARP2 and ARP3) closely resemble the structure of monomeric actin and serve as nucleation sites for new actin filaments. The regulation of actin cytoskeleton rearrangements by ARP2/3 is important for processes like cell locomotion, phagocytosis, and intracellular motility of lipid vesicles, and they have been shown to decrease with age [51]. Other molecules involved in actin cytoskeleton remodeling found in higher amounts in E-ALF were the Ras GTPase-activating like proteins (IQGA1 and IQGA2). These proteins constitute a complex of proteins evolutionarily conserved across species shown to participate in cytoskeleton dynamics, signaling, and cell migration, with several roles identified during microbial pathogenesis by organisms that include important respiratory pathogens [52]. We speculated a potential dysregulation of the RAC signaling pathway, evidenced by lower levels in E-ALF of Coronin 1C (COR1C), with implications in cell migration by regulating the activation and subcellular location of the RAC family of small GTPases, such as RAC2 (**Table 2**). RAC GTPase activities involve secretion, phagocytosis, regulation of superoxide production, and cell polarization [53–58]. Interestingly, RAC2 deficiency is also involved in neutrophil immunodeficiency syndrome [59], potentially driving neutrophilia in the lungs of the elderly. Further supporting this, integrins (ITAL & ITB2) [60] and the proto-oncogene VAV [61] were also lower in E-ALF. This could be further supported by a potential activation of the closely related Rho signaling pathway, evidenced by identified lower levels of septins −2, −7, −9, and −11 [62], Moesin and Redixin [63], and myosin light chains [64]. Another closely related protein, CDC42 (p-value = 0.03, log_2_ fold change = −0.411), did not pass the target fold change for consideration. However, its trend further supports a potential dysregulation of cell polarization and motility, tight junction formation and stability, and formation of actin stress fibers in the lungs of the elderly [65–67] (**Fig. 6**).

**Figure 6.**
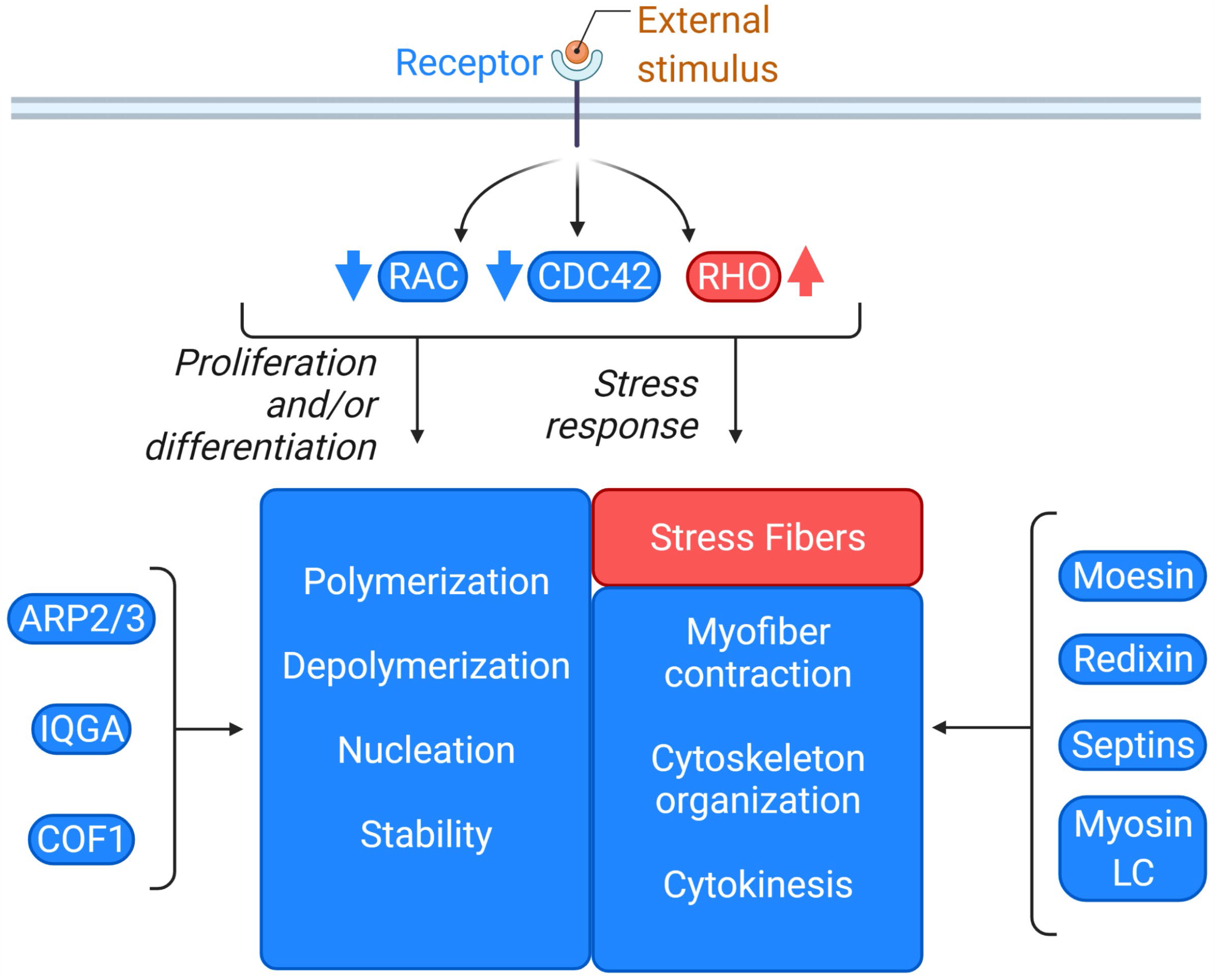
Predicted Actin Cytoskeleton signaling pathway status. The different molecules identified in E-ALF implicated in actin cytoskeleton signaling and possible effects on the lung resident and compartment cells. Blue denotes lower levels of activation, and Red denotes higher levels of activation. This illustration was created with BioRender (https://biorender.com/).

Reduced polymerization of actin in alveolar compartment cells in the elderly would be predicted to directly affect the ability of phagocytes to perform their clearance functions and could impair the timeliness and performance of phagolysosome fusion [68, 69]. Related to actin polymerization, filamin A (FLNA) and its paralog filamin C (FLNC) were also significantly lower in E-ALF. Filamins are proteins that crosslink actin filaments into orthogonal networks in the cytoplasm and actively participate in anchoring membrane proteins for the actin cytoskeleton. Both FLNA and FLNC are implicated in aberrant cell junction organizations, which could affect the optimal lung function in the elderly [70–72].

Other proteins that were highly significantly lower in E-ALF (**Table 2**) were THMS2, a macrophage immunomodulator of B cell responses thought to increase the sensitivity of immature B cells to rare and low-affinity antigens through PLCG2, which was also found significantly lower in the elderly [73, 74]. Caveolin-1 (CAV1), significantly reduced in the ALF of the elderly, is part of the family of the caveolins, necessary for caveolae formation, an endocytic and transcytic clearance and signaling network present in cells of the lung epithelium (e.g., AT-I) and alveolar macrophages, among others [75]. Indeed, downregulation of CAV1 is linked to inflammasome activation [76] and results in different pulmonary disease states, including inflammatory diseases [75]. Related to the formation and stabilization of caveolae in the lung, EH domain-containing protein 2 (EHD2) was also lower in the ALF of the elderly [77], as well as dynamins 1 and 2 (DNM1L and DYN2) [78–80]. Another protein involved in endocytosis of caveolae highly significantly lower in E-ALF, PACN2, is activated by dynamin 1, further indicating dysregulation of caveolae formation, stability, and endocytosis [81, 82]. Coiled-coil domain-containing protein 93 (CCD93), a protein involved in endosome recycling [83], was found to be significantly lower. Chloride intracellular channel 4 (CLIC4) protein was detected at significantly lower levels in E-ALF. CLIC4 regulates fundamental cellular processes, including stabilization of cell membrane potential, transepithelial transport, maintenance of intracellular pH, and regulation of cell volume, which could be directly linked to aberrant membrane cellular transport in the lung cells of elderly individuals [84]. Furthermore, low levels of the Four And A Half LIM domains 1 (FHL1) found in E-ALF could be related to lung function dystrophy [85], and low levels of protein Tyrosine Phosphatase Non-Receptor Type 6 (PTN6) could affect cellular growth and differentiation of migrating cell into the lungs in the elderly [86].

As we described earlier, the soluble surfactant proteins with antimicrobial properties: pulmonary surfactant-associated proteins A and D (SFTPA and SFTPD, respectively), were significantly lower in the elderly ALF, although unexpectedly, SFTPD was not significant (p = 0.056). SFTPA and SFTPD are very important initiators of the innate immune response in the lung, with lower levels associated with susceptibility to several respiratory pathogens in the elderly [13, 14, 87–89]

In this study, we performed a proteomics analysis comparing E-ALF vs. A-ALF composition and we discussed some of the significantly altered proteins and potential pathways implicated. More pathways that were overrepresented in E-ALF are reported in **Table 3**. Our data reported herein should serve as a baseline to understand the biochemical changes that occur in the lung mucosa of the elderly. However, it is clear that due to the difficulty in obtaining samples for these types of studies, and especially the challenge of obtaining samples from individuals of very advanced age (65+), there will need to be a further development of *in vitro* (i.e., 3-dimensional organs, organoids and tissues [90]), and *in vivo* animal models (i.e., aged *Callithrix jacchus [91]* and aged rodents [92, 93]), to study changes in the lung with age. The importance of this understudied area is highlighted by the fact that life expectancy will increase during this century (forecasted to be over 100 years old by year 2,100, in high-income countries) [94].

## MATERIALS AND METHODS

### Human Subjects and Ethics Statement

Human subject studies were carried out in strict accordance with the US Code of Federal and Local Regulations (The Ohio State University IRB numbers: 2008H0135 & 2008H0119 and Texas Biomedical Research Institute/UT-Health San Antonio/ South Texas Veterans Health Care System IRB number: HSC20170667H). Bronchoalveolar lavage fluid (BALF) from healthy adults (aged 23-48 years) and elderly (aged 62-73 years) individuals were recruited from both sexes (adults, female: male ratio of 33%:67%; and elderly, female:male ratio of 50%:50%) immediately prior to partial lung resection, without discrimination of race or ethnicity after informed written consent. For all donors, their preoperative diagnosis was lung nodule, lung nodule with mass, or abnormal parathyroid and thymus, without clinical features. Human donors with these specific comorbidities were excluded: smokers; injection/non-injection drug users; excessive alcohol users; or those with acute pneumonia, upper/lower respiratory tract infections, any kind of acute illness/chronic condition, heart disease, diabetes, asthma, chronic sinusitis/bronchitis, chronic obstructive pulmonary disease, renal failure, liver failure, hepatitis, thyroid disease, rheumatoid arthritis, immunosuppression or taking nonsteroidal anti-inflammatory agents, human immunodeficiency virus (HIV)/AIDS, cancer requiring chemotherapy, leukemia/lymphoma, seizure history, blood disorders, lidocaine allergies (used during the BAL), pregnancy, nontuberculous mycobacterial infections, and TB, as previously described [13].

### BALF processing to obtain ALF

Human bronchoalveolar lavage fluid (BALF) was collected and concentrated to obtain the ALF as previously described [22]. Briefly, BALF was centrifuged to remove cells, the supernatant was sterile-filtered using a 0.22 μm filter, and concentrated ~20X by ultrafiltration with a ten kDa cut-off centrifugal filter (Millipore-Sigma, UFC901008) to obtain a physiological concentration present within the lung as we previously described [13, 14, 22, 24, 87].

### Proteomic Analyses

ALF samples were mixed with 5% SDS/50 mM triethylammonium bicarbonate (TEAB) in the presence of protease and phosphatase inhibitors (Halt; Thermo Scientific). Aliquots corresponding to 50 μg protein (EZQ™ Protein Quantitation Kit; Thermo Fisher) were reduced with tris(2-carboxyethyl) phosphine hydrochloride (TCEP), alkylated in the dark with iodoacetamide and applied to S-Traps (micro; Protifi) for tryptic digestion (sequencing grade; Promega) in 50 mM TEAB. Peptides were eluted from the S-Traps with 0.2% formic acid in 50% aqueous acetonitrile and quantified using Pierce™ Quantitative Fluorometric Peptide Assay (Thermo Scientific).

Data-independent acquisition mass spectrometry was conducted on an Orbitrap Fusion Lumos mass spectrometer (Thermo Scientific). On-line HPLC separation was accomplished with an RSLC NANO HPLC system (Thermo Scientific/Dionex: column, PicoFrit^™^ (New Objective; 75 μm i.d.) packed to 15 cm with C18 adsorbent (Vydac; 218MS 5 μm, 300 Å); mobile phase A, 0.5% acetic acid (HAc)/0.005% trifluoroacetic acid (TFA) in water; mobile phase B, 90% acetonitrile/0.5% HAc/0.005% TFA/9.5% water; gradient 3 to 42% B in 120 min; flow rate, 0.4 μl/min. A pool was made of all of the samples, and 1-μg peptide aliquots were analyzed using gas-phase fractionation and 4-m/z windows (30k resolution for precursor and product ion scans, all in the orbitrap) to create a DIA chromatogram library [95] by searching against a Prosit-generated predicted spectral library [96] based on the UniProt_human_20191022 protein sequence database. Experimental samples were randomized for sample preparation and analysis. Injections of 1 μg of peptides were employed. MS data for experimental samples were acquired in the orbitrap using 8-m/z windows (staggered; 30k resolution for precursor and product ion scans) and searched against the chromatogram library. Scaffold DIA (v3.0.0; Proteome Software) was used for all DIA data processing.

### Data analyses

DEPs were selected based on the following thresholds, Log_2_ Fold Change < −0.5 or > 0.5, p-value < 0.055 as given by Benjamini Hochberg multiple testing correction of False Discovery Rate set at 20%, a liberal threshold to allow for the maximum discovery of DEPs. Microsoft Excel 2019 (Microsoft Corporation) was used for the generation and handling of tables. Plotly Dash Chart [97] was used to generate bar and bubble charts. ClustVis [98] was used for the creation of hierarchical clustering heat maps. DEPs were annotated using the 2021-08-18 release of Human GOA (Gene Ontology Consortium). Each DEP was assigned all GO annotations for what a match existed. Each group of annotations was counted to perform GO annotation classifications. Ingenuity Pathway Analysis (v65367011; QIAGEN) was used to perform pathway analysis and regulator prediction analyses.

## AUTHOR CONTRIBUTIONS

A.G-V. analyzed the data. A.G-V., A.M.O-F., J.I.M., and A.A-G. processed samples. H.S., R.E.M., D.M-C., and J.P. procured samples. Y.W. critically revised the data analysis. S.T.W. supervised MS analysis and performed the primary analysis of DIA-MS data. A.G-V. and J.B.T. conceptualized the study, designed the experiments, and wrote the manuscript. Y.W., S.T.W., L.S.S., J.T., and J.B.T. critically reviewed the manuscript. L.S.S., J.T., and J.B.T. provided funding. All authors read and approved the final version of the manuscript.

## ACKNOWLEDGEMENTS

Mass spectrometry analyses were conducted in the Institutional Mass Spectrometry Laboratory of the University of Texas Health Science Center at San Antonio (UTHSCSA), with support from UTHSCSA for the laboratory and the University of Texas System for purchase of the Orbitrap Fusion Lumos mass spectrometer; expert technical assistance of Mr. Sammy Pardo and Ms. Dana Molleur is gratefully acknowledged. The authors would also like to thank Drs. Colwyn Headley and Tucker Piergallini for their assistance with IPA analysis.

## CONFLICT OF INTEREST

The authors declare no conflict of interest exists.

## FUNDING

This study was supported by the National Institute on Aging (NIA), National Institutes of Health (NIH) (Grant number P01 AG-051428 to J.T., J.B.T., B.I.R, and L.S.S.), and J.B.T. was partially supported by Robert J. Kleberg, Jr. and Helen C. Kleberg Foundation. The content is solely the responsibility of the authors and does not necessarily represent the official views of the National Institutes of Health. A.M.O-F. was supported by the Douglass Graduate Fellowship at Texas Biomedical Research Institute.

## SUPPLEMENTAL MATERIAL

**Supplemental Table S1. Canonical pathways.** A table with all (417) significantly enriched Canonical Pathways identified by IPA analysis, showing −log(p-value) of the protein overlap, a ratio of DEPs in dataset belonging to pathway / Proteins in pathway for reference dataset, and an activation z-score, an estimation of the overall activation of the pathway based on the status of the molecules identified in E-ALF.

**Supplemental Fig S1. Gene Ontology annotations, distribution of all proteins.** Stacked bar chart of the total number of proteins identified by DIA-MS (Blue [Lower] and Red [Higher]) belonging to the different GO classifications (Biological Process, Cellular Component, and Molecular Function).

## Notes

### Competing Interest Statement

The authors have declared no competing interest.

