## Supplemental Table 1 for "The aging lung mucosa: A proteomics study"

| Ingenuity Canonical Pathways | -log(p-value) | Ratio | z-score |
|---|---|---|---|
| Actin Cytoskeleton Signaling | 22.8 | 0.155 | -4.004 |
| Remodeling of Epithelial Adherens Junctions | 20.8 | 0.324 | -2.828 |
| Integrin Signaling | 15.9 | 0.136 | -3.024 |
| Signaling by Rho Family GTPases | 14.9 | 0.116 | -4.6 |
| RHOGDI Signaling | 14.9 | 0.13 | 3.273 |
| Leukocyte Extravasation Signaling | 14.2 | 0.135 | -2.858 |
| RHOA Signaling | 13.6 | 0.169 | -4.146 |
| Regulation of Actin-based Motility by Rho | 13.1 | 0.172 | -3.153 |
| Axonal Guidance Signaling | 12.7 | 0.0771 | NaN |
| RAC Signaling | 12.6 | 0.152 | -3.3 |
| Clathrin-mediated Endocytosis Signaling | 11.5 | 0.119 | NaN |
| Germ Cell-Sertoli Cell Junction Signaling | 10.8 | 0.123 | NaN |
| Fcγ Receptor-mediated Phagocytosis in Macrophages and Monocytes | 10.5 | 0.17 | -3 |
| EIF2 Signaling | 10.2 | 0.103 | 1.387 |
| Phagosome Maturation | 8.76 | 0.113 | NaN |
| Actin Nucleation by ARP-WASP Complex | 8.56 | 0.151 | -3.162 |
| Sertoli Cell-Sertoli Cell Junction Signaling | 8.48 | 0.0971 | NaN |
| VEGF Signaling | 8.2 | 0.141 | -2.121 |
| Epithelial Adherens Junction Signaling | 8.01 | 0.108 | -1.213 |
| Ephrin Receptor Signaling | 7.9 | 0.0945 | -3.638 |
| IL-8 Signaling | 7.55 | 0.09 | -1.886 |
| Production of Nitric Oxide and Reactive Oxygen Species in Macrophages | 7.49 | 0.0942 | 0.258 |
| Semaphorin Neuronal Repulsive Signaling Pathway | 7.44 | 0.106 | 0 |
| Synaptogenesis Signaling Pathway | 7.37 | 0.0737 | -2.4 |
| FAK Signaling | 7.25 | 0.12 | NaN |
| UDP-N-acetyl-D-galactosamine Biosynthesis II | 7.14 | 0.462 | -0.816 |
| 14-3-3-mediated Signaling | 6.8 | 0.11 | -1.89 |
| Agranulocyte Adhesion and Diapedesis | 6.74 | 0.0841 | NaN |
| fMLP Signaling in Neutrophils | 6.63 | 0.107 | -3.464 |
| Gap Junction Signaling | 6.52 | 0.0859 | NaN |
| Role of Tissue Factor in Cancer | 6.44 | 0.112 | NaN |
| Virus Entry via Endocytic Pathways | 6.13 | 0.115 | NaN |
| Paxillin Signaling | 5.95 | 0.111 | -1.897 |
| Phagosome Formation | 5.81 | 0.0479 | -4.004 |
| PFKFB4 Signaling Pathway | 5.63 | 0.174 | -0.707 |
| LXR/RXR Activation | 5.35 | 0.0976 | 2.111 |
| Dilated Cardiomyopathy Signaling Pathway | 5.3 | 0.089 | 0.905 |
| ILK Signaling | 5.16 | 0.0758 | -1.941 |
| G Beta Gamma Signaling | 5.14 | 0.093 | -2.309 |
| tRNA Charging | 5.09 | 0.179 | -2.646 |
| Regulation of eIF4 and p70S6K Signaling | 5.01 | 0.0782 | NaN |
| Caveolar-mediated Endocytosis Signaling | 4.88 | 0.12 | NaN |
| PAK Signaling | 4.76 | 0.0932 | -2.121 |

### Supplemental Table 1, continued

| Ingenuity Canonical Pathways | -log(p-value) | Ratio | z-score |
|---|---|---|---|
| Renin-Angiotensin Signaling | 4.7 | 0.0917 | -2.111 |
| Coronavirus Replication Pathway | 4.66 | 0.156 | 2.646 |
| FXR/RXR Activation | 4.5 | 0.0873 | NaN |
| Reelin Signaling in Neurons | 4.5 | 0.0873 | -2.53 |
| Cardiac Hypertrophy Signaling | 4.38 | 0.062 | -1.941 |
| Acute Phase Response Signaling | 4.21 | 0.0703 | 0.333 |
| IL-12 Signaling and Production in Macrophages | 4.19 | 0.0809 | NaN |
| ERK/MAPK Signaling | 4.15 | 0.0654 | -2.111 |
| CXCR4 Signaling | 4.02 | 0.0719 | -2.714 |
| Estrogen Receptor Signaling | 3.93 | 0.0495 | -1.414 |
| Mechanisms of Viral Exit from Host Cells | 3.91 | 0.146 | NaN |
| Natural Killer Cell Signaling | 3.89 | 0.0653 | -1.387 |
| Cellular Effects of Sildenafil (Viagra) | 3.84 | 0.0738 | NaN |
| Tight Junction Signaling | 3.76 | 0.0674 | NaN |
| Semaphorin Signaling in Neurons | 3.75 | 0.113 | NaN |
| HIF1α Signaling | 3.7 | 0.0625 | -1.732 |
| Atherosclerosis Signaling | 3.68 | 0.0769 | NaN |
| Thrombin Signaling | 3.66 | 0.0619 | -3 |
| Trehalose Degradation II (Trehalase) | 3.64 | 0.429 | NaN |
| Granulocyte Adhesion and Diapedesis | 3.52 | 0.0635 | NaN |
| Regulation of Cellular Mechanics by Calpain Protease | 3.51 | 0.0899 | 0 |
| Agrin Interactions at Neuromuscular Junction | 3.42 | 0.1 | -1.134 |
| ERBB Signaling | 3.35 | 0.0851 | -2.121 |
| Ephrin B Signaling | 3.34 | 0.0972 | -0.816 |
| Glioma Invasiveness Signaling | 3.31 | 0.0959 | -1.633 |
| Coagulation System | 3.29 | 0.143 | 1.342 |
| Colorectal Cancer Metastasis Signaling | 3.13 | 0.052 | -0.302 |
| Pentose Phosphate Pathway | 3.12 | 0.3 | NaN |
| GP6 Signaling Pathway | 3.11 | 0.0709 | -0.707 |
| Protein Kinase A Signaling | 3.08 | 0.0448 | 0.277 |
| Apoptosis Signaling | 3.05 | 0.0769 | 1.414 |
| Calcium Signaling | 3 | 0.0556 | NaN |
| α-Adrenergic Signaling | 3 | 0.0755 | -2.236 |
| p70S6K Signaling | 2.99 | 0.0682 | -2.449 |
| GDP-glucose Biosynthesis | 2.99 | 0.273 | NaN |
| GNRH Signaling | 2.98 | 0.0585 | -1.897 |
| CCR3 Signaling in Eosinophils | 2.92 | 0.0667 | -1.633 |
| Glycolysis I | 2.91 | 0.16 | 0 |
| Gluconeogenesis I | 2.91 | 0.16 | 0 |
| PPARα/RXRα Activation | 2.89 | 0.057 | 1.897 |
| Hepatic Fibrosis Signaling Pathway | 2.88 | 0.043 | -2.84 |
| Apelin Adipocyte Signaling Pathway | 2.88 | 0.0814 | -0.816 |
| Glucose and Glucose-1-phosphate Degradation | 2.87 | 0.25 | NaN |

### Supplemental Table 1, continued

| Ingenuity Canonical Pathways | -log(p-value) | Ratio | z-score |
|---|---|---|---|
| Role of MAPK Signaling in Promoting the Pathogenesis of Influenza | 2.84 | 0.0714 | -2.828 |
| Iron homeostasis signaling pathway | 2.83 | 0.0647 | NaN |
| Amyotrophic Lateral Sclerosis Signaling | 2.77 | 0.0696 | 0 |
| Fc Epsilon RI Signaling | 2.7 | 0.0678 | -2.828 |
| Airway Pathology in Chronic Obstructive Pulmonary Disease | 2.7 | 0.0678 | NaN |
| Cholecystokinin/Gastrin-mediated Signaling | 2.68 | 0.0672 | -2.646 |
| Pentose Phosphate Pathway (Oxidative Branch) | 2.67 | 0.5 | NaN |
| N-acetylglucosamine Degradation II | 2.67 | 0.5 | NaN |
| Role of IL-17A in Psoriasis | 2.66 | 0.214 | NaN |
| Colanic Acid Building Blocks Biosynthesis | 2.66 | 0.214 | NaN |
| Melanocyte Development and Pigmentation Signaling | 2.66 | 0.0745 | -1.134 |
| Corticotropin Releasing Hormone Signaling | 2.63 | 0.0604 | -1.414 |
| Molecular Mechanisms of Cancer | 2.6 | 0.0404 | NaN |
| Melatonin Signaling | 2.59 | 0.0833 | -1.633 |
| Amyloid Processing | 2.54 | 0.098 | NaN |
| Apelin Cardiomyocyte Signaling Pathway | 2.53 | 0.0707 | -2.646 |
| Extrinsic Prothrombin Activation Pathway | 2.49 | 0.188 | NaN |
| Macropinocytosis Signaling | 2.47 | 0.0789 | -1 |
| Airway Inflammation in Asthma | 2.46 | 0.121 | NaN |
| CDC42 Signaling | 2.41 | 0.0365 | -3.771 |
| IGF-1 Signaling | 2.41 | 0.0673 | -1.342 |
| HGF Signaling | 2.4 | 0.0606 | -1.633 |
| IL-3 Signaling | 2.39 | 0.0759 | -0.816 |
| Chemokine Signaling | 2.36 | 0.075 | -1.633 |
| Role of PKR in Interferon Induction and Antiviral Response | 2.32 | 0.0588 | -1.633 |
| Urea Cycle | 2.28 | 0.333 | NaN |
| UDP-N-acetyl-D-glucosamine Biosynthesis II | 2.28 | 0.333 | NaN |
| Rapoport-Luebering Glycolytic Shunt | 2.28 | 0.333 | NaN |
| VEGF Family Ligand-Receptor Interactions | 2.26 | 0.0714 | -2.236 |
| Gαq Signaling | 2.25 | 0.0529 | -1.633 |
| Coronavirus Pathogenesis Pathway | 2.22 | 0.0493 | -3.162 |
| Prolactin Signaling | 2.21 | 0.0698 | -2.236 |
| Inhibition of Matrix Metalloproteases | 2.19 | 0.103 | -1 |
| BMP signaling pathway | 2.18 | 0.069 | -1.633 |
| Thrombopoietin Signaling | 2.15 | 0.0794 | -2.236 |
| AMPK Signaling | 2.14 | 0.0455 | 0.378 |
| Neuregulin Signaling | 2.13 | 0.0598 | -2.236 |
| Sphingosine-1-phosphate Signaling | 2.11 | 0.0593 | -1.342 |
| NGF Signaling | 2.11 | 0.0593 | -2.646 |
| Intrinsic Prothrombin Activation Pathway | 2.08 | 0.0952 | 2 |
| IL-1 Signaling | 2.03 | 0.0638 | 0 |
| Endocannabinoid Developing Neuron Pathway | 2.02 | 0.0569 | -2.236 |
| ERBB4 Signaling | 2.01 | 0.0735 | -2.236 |

### Supplemental Table 1, continued

| Ingenuity Canonical Pathways | -log(p-value) | Ratio | z-score |
|---|---|---|---|
| Role of NFAT in Cardiac Hypertrophy | 1.99 | 0.0455 | -1.897 |
| Tumoricidal Function of Hepatic Natural Killer Cells | 1.98 | 0.125 | NaN |
| Salvage Pathways of Pyrimidine Ribonucleotides | 1.94 | 0.0612 | -0.816 |
| Endothelin-1 Signaling | 1.94 | 0.0471 | -2.333 |
| Salvage Pathways of Pyrimidine Deoxyribonucleotides | 1.92 | 0.222 | NaN |
| P2Y Purigenic Receptor Signaling Pathway | 1.91 | 0.0543 | -1.134 |
| RAR Activation | 1.87 | 0.0459 | NaN |
| Leptin Signaling in Obesity | 1.86 | 0.0676 | NaN |
| Gα12/13 Signaling | 1.84 | 0.0526 | -2.646 |
| Insulin Secretion Signaling Pathway | 1.83 | 0.041 | -1.897 |
| Adrenomedullin signaling pathway | 1.83 | 0.0452 | -2.333 |
| Osteoarthritis Pathway | 1.82 | 0.0427 | 1.342 |
| MYC Mediated Apoptosis Signaling | 1.82 | 0.08 | -1 |
| UVC-Induced MAPK Signaling | 1.79 | 0.0784 | -2 |
| NRF2-mediated Oxidative Stress Response | 1.78 | 0.0422 | -2 |
| Maturity Onset Diabetes of Young (MODY) Signaling | 1.77 | 0.0641 | NaN |
| Gαi Signaling | 1.76 | 0.0507 | -1.342 |
| MSP-RON Signaling In Cancer Cells Pathway | 1.76 | 0.0507 | -1.89 |
| Glucocorticoid Receptor Signaling | 1.76 | 0.0327 | NaN |
| Opioid Signaling Pathway | 1.74 | 0.0399 | -1.667 |
| Renal Cell Carcinoma Signaling | 1.73 | 0.0625 | -1.342 |
| Melatonin Degradation III | 1.72 | 1 | NaN |
| Glutamine Biosynthesis I | 1.72 | 1 | NaN |
| UDP-N-acetyl-D-galactosamine Biosynthesis I | 1.72 | 1 | NaN |
| PI3K Signaling in B Lymphocytes | 1.69 | 0.049 | -2.236 |
| Endocannabinoid Cancer Inhibition Pathway | 1.69 | 0.049 | -1.134 |
| mTOR Signaling | 1.67 | 0.0425 | -1.342 |
| Tumor Microenvironment Pathway | 1.65 | 0.0447 | 0 |
| FcγRIIB Signaling in B Lymphocytes | 1.63 | 0.0588 | NaN |
| LPS-stimulated MAPK Signaling | 1.63 | 0.0588 | -1.342 |
| GPCR-Mediated Nutrient Sensing in Enteroendocrine Cells | 1.61 | 0.0517 | -0.816 |
| PDGF Signaling | 1.61 | 0.0581 | -1.342 |
| Xenobiotic Metabolism AHR Signaling Pathway | 1.59 | 0.0575 | 0.447 |
| Neuroprotective Role of THOP1 in Alzheimer's Disease | 1.58 | 0.0508 | 0.447 |
| Inhibition of Angiogenesis by TSP1 | 1.57 | 0.0882 | NaN |
| Nitric Oxide Signaling in the Cardiovascular System | 1.57 | 0.0504 | -0.447 |
| Endometrial Cancer Signaling | 1.55 | 0.0667 | -2 |
| Leukotriene Biosynthesis | 1.54 | 0.143 | NaN |
| Crosstalk between Dendritic Cells and Natural Killer Cells | 1.51 | 0.0549 | NaN |
| Necroptosis Signaling Pathway | 1.5 | 0.0446 | -1.134 |
| D-myo-inositol-5-phosphate Metabolism | 1.49 | 0.0417 | -0.707 |
| Superpathway of Citrulline Metabolism | 1.49 | 0.133 | NaN |
| Ovarian Cancer Signaling | 1.48 | 0.0443 | -0.816 |

### Supplemental Table 1, continued

| Ingenuity Canonical Pathways | -log(p-value) | Ratio | z-score |
|---|---|---|---|
| eNOS Signaling | 1.47 | 0.044 | -1.134 |
| Granzyme B Signaling | 1.43 | 0.125 | NaN |
| Adenosine Nucleotides Degradation II | 1.43 | 0.125 | NaN |
| Death Receptor Signaling | 1.43 | 0.0521 | -1.342 |
| Pyridoxal 5'-phosphate Salvage Pathway | 1.42 | 0.0606 | -1 |
| Palmitate Biosynthesis I (Animals) | 1.42 | 0.5 | NaN |
| L-cysteine Degradation III | 1.42 | 0.5 | NaN |
| Sulfate Activation for Sulfonation | 1.42 | 0.5 | NaN |
| Fatty Acid Biosynthesis Initiation II | 1.42 | 0.5 | NaN |
| GDP-L-fucose Biosynthesis I (from GDP-D-mannose) | 1.42 | 0.5 | NaN |
| Synaptic Long Term Potentiation | 1.42 | 0.0465 | -1.633 |
| Histamine Degradation | 1.39 | 0.118 | NaN |
| Glioblastoma Multiforme Signaling | 1.33 | 0.0409 | -2.449 |
| Oncostatin M Signaling | 1.31 | 0.0698 | NaN |
| B Cell Development | 1.31 | 0.0698 | NaN |
| ERK5 Signaling | 1.31 | 0.0556 | -1 |
| Netrin Signaling | 1.31 | 0.0556 | -1 |
| Granzyme A Signaling | 1.3 | 0.105 | NaN |
| Purine Nucleotides Degradation II (Aerobic) | 1.3 | 0.105 | NaN |
| GPCR-Mediated Integration of Enteroendocrine Signaling Exemplified by an L Cell | 1.29 | 0.0548 | 1 |
| G Protein Signaling Mediated by Tubby | 1.28 | 0.0682 | NaN |
| Insulin Receptor Signaling | 1.28 | 0.0429 | -1.633 |
| D-myo-inositol (1,4,5,6)-Tetrakisphosphate Biosynthesis | 1.27 | 0.0398 | -0.378 |
| D-myo-inositol (3,4,5,6)-tetrakisphosphate Biosynthesis | 1.27 | 0.0398 | -0.378 |
| Inflammasome pathway | 1.26 | 0.1 | NaN |
| Ascorbate Recycling (Cytosolic) | 1.25 | 0.333 | NaN |
| Glycerol-3-phosphate Shuttle | 1.25 | 0.333 | NaN |
| Glutamate Degradation II | 1.25 | 0.333 | NaN |
| N-acetylglucosamine Degradation I | 1.25 | 0.333 | NaN |
| Aspartate Biosynthesis | 1.25 | 0.333 | NaN |
| Xenobiotic Metabolism General Signaling Pathway | 1.24 | 0.042 | -1.633 |
| GDNF Family Ligand-Receptor Interactions | 1.24 | 0.0526 | -2 |
| TREM1 Signaling | 1.22 | 0.0519 | -1 |
| Angiopoietin Signaling | 1.22 | 0.0519 | -1 |
| nNOS Signaling in Neurons | 1.21 | 0.0638 | NaN |
| Ephrin A Signaling | 1.21 | 0.0638 | NaN |
| IL-7 Signaling Pathway | 1.2 | 0.0513 | 0 |
| Endocannabinoid Neuronal Synapse Pathway | 1.19 | 0.0408 | -1.633 |
| CDK5 Signaling | 1.19 | 0.0446 | -0.447 |
| Breast Cancer Regulation by Stathmin1 | 1.19 | 0.0287 | -2.668 |
| FLT3 Signaling in Hematopoietic Progenitor Cells | 1.17 | 0.05 | -2 |
| PTEN Signaling | 1.16 | 0.04 | 2 |
| Role of MAPK Signaling in the Pathogenesis of Influenza | 1.15 | 0.0494 | NaN |

### Supplemental Table 1, continued

| Ingenuity Canonical Pathways | -log(p-value) | Ratio | z-score |
|---|---|---|---|
| 3-phosphoinositide Degradation | 1.15 | 0.0372 | -0.378 |
| Cell Cycle: G2/M DNA Damage Checkpoint Regulation | 1.15 | 0.06 | NaN |
| Differential Regulation of Cytokine Production in Intestinal Epithelial Cells by IL-17 | 1.15 | 0.087 | NaN |
| Vitamin-C Transport | 1.15 | 0.087 | NaN |
| Bladder Cancer Signaling | 1.14 | 0.0431 | NaN |
| JAK/STAT Signaling | 1.14 | 0.0488 | -1 |
| CD28 Signaling in T Helper Cells | 1.13 | 0.029 | -2.138 |
| Relaxin Signaling | 1.13 | 0.0392 | 0 |
| Heme Degradation | 1.13 | 0.25 | NaN |
| Arginine Degradation I (Arginase Pathway) | 1.13 | 0.25 | NaN |
| Oxidized GTP and dGTP Detoxification | 1.13 | 0.25 | NaN |
| L-cysteine Degradation I | 1.13 | 0.25 | NaN |
| MSP-RON Signaling In Macrophages Pathway | 1.13 | 0.0427 | -0.447 |
| Xenobiotic Metabolism PXR Signaling Pathway | 1.12 | 0.0366 | 0.378 |
| TNFR1 Signaling | 1.11 | 0.0577 | NaN |
| UVB-Induced MAPK Signaling | 1.11 | 0.0577 | NaN |
| FGF Signaling | 1.11 | 0.0476 | -1 |
| Superpathway of Inositol Phosphate Compounds | 1.1 | 0.0345 | -0.707 |
| Synaptic Long Term Depression | 1.09 | 0.0361 | -2.646 |
| Hepatic Fibrosis / Hepatic Stellate Cell Activation | 1.09 | 0.0361 | NaN |
| HIPPO signaling | 1.09 | 0.0471 | NaN |
| Gαs Signaling | 1.08 | 0.0413 | NaN |
| Th1 Pathway | 1.07 | 0.041 | NaN |
| IL-17A Signaling in Gastric Cells | 1.06 | 0.0769 | NaN |
| Primary Immunodeficiency Signaling | 1.05 | 0.0545 | NaN |
| EGF Signaling | 1.05 | 0.0545 | NaN |
| PI3K/AKT Signaling | 1.05 | 0.0352 | -1.342 |
| Glioma Signaling | 1.04 | 0.0403 | -2.236 |
| Arsenate Detoxification I (Glutaredoxin) | 1.04 | 0.2 | NaN |
| Creatine-phosphate Biosynthesis | 1.04 | 0.2 | NaN |
| Serine Biosynthesis | 1.04 | 0.2 | NaN |
| CMP-N-acetylneuraminate Biosynthesis I (Eukaryotes) | 1.04 | 0.2 | NaN |
| Myo-inositol Biosynthesis | 1.04 | 0.2 | NaN |
| Citrulline-Nitric Oxide Cycle | 1.04 | 0.2 | NaN |
| Galactose Degradation I (Leloir Pathway) | 1.04 | 0.2 | NaN |
| Huntington's Disease Signaling | 1.03 | 0.032 | 0.816 |
| NAD Salvage Pathway II | 1.03 | 0.0741 | NaN |
| Unfolded protein response | 1.02 | 0.0444 | NaN |
| 3-phosphoinositide Biosynthesis | 1.02 | 0.0347 | -0.378 |
| Ferroptosis Signaling Pathway | 1.02 | 0.0397 | -1.342 |
| CNTF Signaling | 1.02 | 0.0526 | NaN |
| Role of IL-17A in Arthritis | 1.02 | 0.0526 | NaN |
| MSP-RON Signaling Pathway | 1 | 0.0517 | NaN |

### Supplemental Table 1, continued

| Ingenuity Canonical Pathways | -log(p-value) | Ratio | z-score |
|---|---|---|---|
| IL-4 Signaling | 0.987 | 0.043 | NaN |
| Sonic Hedgehog Signaling | 0.975 | 0.069 | NaN |
| Androgen Signaling | 0.971 | 0.0355 | -1 |
| SPINK1 Pancreatic Cancer Pathway | 0.967 | 0.05 | NaN |
| Arginine Biosynthesis IV | 0.959 | 0.167 | NaN |
| Arginine Degradation VI (Arginase 2 Pathway) | 0.959 | 0.167 | NaN |
| Pentose Phosphate Pathway (Non-oxidative Branch) | 0.959 | 0.167 | NaN |
| Glycerol Degradation I | 0.959 | 0.167 | NaN |
| Selenocysteine Biosynthesis II (Archaea and Eukaryotes) | 0.959 | 0.167 | NaN |
| Tryptophan Degradation to 2-amino-3-carboxymuconate Semialdehyde | 0.959 | 0.167 | NaN |
| GDP-mannose Biosynthesis | 0.959 | 0.167 | NaN |
| IL-2 Signaling | 0.951 | 0.0492 | NaN |
| TGF-β Signaling | 0.947 | 0.0417 | -2 |
| Sperm Motility | 0.932 | 0.0315 | -1 |
| p53 Signaling | 0.924 | 0.0408 | NaN |
| White Adipose Tissue Browning Pathway | 0.917 | 0.0368 | -0.447 |
| Glutathione-mediated Detoxification | 0.903 | 0.0625 | NaN |
| Ethanol Degradation II | 0.903 | 0.0625 | NaN |
| Erythropoietin Signaling Pathway | 0.9 | 0.0339 | -1.633 |
| Superpathway of Serine and Glycine Biosynthesis I | 0.896 | 0.143 | NaN |
| Adenine and Adenosine Salvage III | 0.896 | 0.143 | NaN |
| Ceramide Degradation | 0.896 | 0.143 | NaN |
| Aspartate Degradation II | 0.896 | 0.143 | NaN |
| Neuropathic Pain Signaling In Dorsal Horn Neurons | 0.889 | 0.0396 | -1 |
| ERB2-ERBB3 Signaling | 0.889 | 0.0462 | NaN |
| Apelin Endothelial Signaling Pathway | 0.889 | 0.036 | -2.236 |
| Dopamine-DARPP32 Feedback in cAMP Signaling | 0.87 | 0.0331 | -0.447 |
| IL-17A Signaling in Airway Cells | 0.863 | 0.0448 | NaN |
| 4-1BB Signaling in T Lymphocytes | 0.86 | 0.0588 | NaN |
| Mouse Embryonic Stem Cell Pluripotency | 0.857 | 0.0385 | -2 |
| Purine Ribonucleosides Degradation to Ribose-1-phosphate | 0.845 | 0.125 | NaN |
| Noradrenaline and Adrenaline Degradation | 0.842 | 0.0571 | NaN |
| Retinoate Biosynthesis I | 0.821 | 0.0556 | NaN |
| Interferon Signaling | 0.821 | 0.0556 | NaN |
| GM-CSF Signaling | 0.821 | 0.0429 | NaN |
| Neuroinflammation Signaling Pathway | 0.815 | 0.0286 | -1.89 |
| Xenobiotic Metabolism CAR Signaling Pathway | 0.815 | 0.0319 | 0.816 |
| Cardiac Hypertrophy Signaling (Enhanced) | 0.812 | 0.026 | -2.714 |
| Growth Hormone Signaling | 0.81 | 0.0423 | NaN |
| Complement System | 0.801 | 0.0541 | NaN |
| Sphingosine and Sphingosine-1-phosphate Metabolism | 0.796 | 0.111 | NaN |
| Citrulline Biosynthesis | 0.796 | 0.111 | NaN |
| Antioxidant Action of Vitamin C | 0.785 | 0.036 | NaN |

### Supplemental Table 1, continued

| Ingenuity Canonical Pathways | -log(p-value) | Ratio | z-score |
|---|---|---|---|
| Docosahexaenoic Acid (DHA) Signaling | 0.783 | 0.0526 | NaN |
| IL-17A Signaling in Fibroblasts | 0.783 | 0.0526 | NaN |
| Neurotrophin/TRK Signaling | 0.747 | 0.0395 | NaN |
| Role of MAPK Signaling in Inhibiting the Pathogenesis of Influenza | 0.747 | 0.0395 | NaN |
| Antiproliferative Role of Somatostatin Receptor 2 | 0.738 | 0.039 | NaN |
| Aryl Hydrocarbon Receptor Signaling | 0.721 | 0.0314 | NaN |
| Xenobiotic Metabolism Signaling | 0.721 | 0.0278 | NaN |
| Purine Nucleotides De Novo Biosynthesis II | 0.717 | 0.0909 | NaN |
| Inhibition of ARE-Mediated mRNA Degradation Pathway | 0.706 | 0.0311 | -0.447 |
| HOTAIR Regulatory Pathway | 0.69 | 0.0307 | 1.342 |
| Aldosterone Signaling in Epithelial Cells | 0.684 | 0.0305 | NaN |
| NAD biosynthesis II (from tryptophan) | 0.684 | 0.0833 | NaN |
| Glycogen Degradation II | 0.684 | 0.0833 | NaN |
| LPS/IL-1 Mediated Inhibition of RXR Function | 0.682 | 0.0279 | NaN |
| Senescence Pathway | 0.674 | 0.0269 | 0.378 |
| Role of IL-17F in Allergic Inflammatory Airway Diseases | 0.672 | 0.0444 | NaN |
| HMGB1 Signaling | 0.664 | 0.0299 | -2 |
| TR/RXR Activation | 0.662 | 0.0357 | NaN |
| PEDF Signaling | 0.662 | 0.0357 | NaN |
| Apelin Pancreas Signaling Pathway | 0.658 | 0.0435 | NaN |
| Acyl-CoA Hydrolysis | 0.654 | 0.0769 | NaN |
| NAD Phosphorylation and Dephosphorylation | 0.654 | 0.0769 | NaN |
| Guanosine Nucleotides Degradation III | 0.654 | 0.0769 | NaN |
| Pancreatic Adenocarcinoma Signaling | 0.652 | 0.0317 | 0 |
| Phospholipase C Signaling | 0.65 | 0.0237 | -2.714 |
| Role of Hypercytokinemia/hyperchemokinemia in the Pathogenesis of Influenza | 0.642 | 0.0349 | NaN |
| IL-6 Signaling | 0.636 | 0.0312 | -2 |
| Apelin Muscle Signaling Pathway | 0.631 | 0.0417 | NaN |
| Th1 and Th2 Activation Pathway | 0.629 | 0.0291 | NaN |
| Glycogen Degradation III | 0.625 | 0.0714 | NaN |
| Urate Biosynthesis/Inosine 5'-phosphate Degradation | 0.625 | 0.0714 | NaN |
| Phenylalanine Degradation IV (Mammalian, via Side Chain) | 0.625 | 0.0714 | NaN |
| Cardiac β-adrenergic Signaling | 0.616 | 0.0287 | NaN |
| Melanoma Signaling | 0.604 | 0.04 | NaN |
| Choline Biosynthesis III | 0.599 | 0.0667 | NaN |
| Acute Myeloid Leukemia Signaling | 0.597 | 0.033 | NaN |
| STAT3 Pathway | 0.585 | 0.0296 | -1 |
| Chondroitin Sulfate Degradation (Metazoa) | 0.575 | 0.0625 | NaN |
| Parkinson's Signaling | 0.575 | 0.0625 | NaN |
| Th2 Pathway | 0.572 | 0.0292 | NaN |
| Non-Small Cell Lung Cancer Signaling | 0.57 | 0.0319 | NaN |
| Stearate Biosynthesis I (Animals) | 0.569 | 0.0377 | NaN |
| Phototransduction Pathway | 0.558 | 0.037 | NaN |

### Supplemental Table 1, continued

| **Ingenuity Canonical Pathways** | **-log(p-value)** | **Ratio** | **z-score** |
|---|---|---|---|
| RAN Signaling | 0.553 | 0.0588 | NaN |
| Dermatan Sulfate Degradation (Metazoa) | 0.553 | 0.0588 | NaN |
| ATM Signaling | 0.547 | 0.0309 | NaN |
| IL-17 Signaling | 0.539 | 0.0267 | 0.447 |
| UVA-Induced MAPK Signaling | 0.538 | 0.0306 | NaN |
| CSDE1 Signaling Pathway | 0.536 | 0.0357 | NaN |
| Differential Regulation of Cytokine Production in Macrophages and T Helper Cells b | 0.532 | 0.0556 | NaN |
| CD27 Signaling in Lymphocytes | 0.526 | 0.0351 | NaN |
| Cancer Drug Resistance By Drug Efflux | 0.516 | 0.0345 | NaN |
| Regulation Of The Epithelial Mesenchymal Transition By Growth Factors Pathway | 0.511 | 0.026 | -1.342 |
| Sumoylation Pathway | 0.5 | 0.0291 | NaN |
| TEC Kinase Signaling | 0.496 | 0.0225 | -2.121 |
| Methylglyoxal Degradation III | 0.493 | 0.05 | NaN |
| Fatty Acid α-oxidation | 0.493 | 0.05 | NaN |
| Putrescine Degradation III | 0.476 | 0.0476 | NaN |
| Endoplasmic Reticulum Stress Pathway | 0.476 | 0.0476 | NaN |
| Chronic Myeloid Leukemia Signaling | 0.472 | 0.028 | NaN |
| Telomerase Signaling | 0.472 | 0.028 | NaN |
| PPAR Signaling | 0.472 | 0.028 | NaN |
| Systemic Lupus Erythematosus In T Cell Signaling Pathway | 0.466 | 0.0219 | -2.138 |
| Pyrimidine Deoxyribonucleotides De Novo Biosynthesis I | 0.46 | 0.0455 | NaN |
| Role of Pattern Recognition Receptors in Recognition of Bacteria and Viruses | 0.456 | 0.0256 | -1 |
| PXR/RXR Activation | 0.45 | 0.0308 | NaN |
| Induction of Apoptosis by HIV1 | 0.45 | 0.0308 | NaN |
| Ethanol Degradation IV | 0.445 | 0.0435 | NaN |
| Role of PI3K/AKT Signaling in the Pathogenesis of Influenza | 0.442 | 0.0303 | NaN |
| Phospholipases | 0.442 | 0.0303 | NaN |
| Prostate Cancer Signaling | 0.44 | 0.0268 | NaN |
| CD40 Signaling | 0.434 | 0.0299 | NaN |
| Serotonin Degradation | 0.434 | 0.0299 | NaN |
| Eicosanoid Signaling | 0.434 | 0.0299 | NaN |
| IL-22 Signaling | 0.431 | 0.0417 | NaN |
| TCA Cycle II (Eukaryotic) | 0.431 | 0.0417 | NaN |
| Tryptophan Degradation III (Eukaryotic) | 0.431 | 0.0417 | NaN |
| Role of JAK1 and JAK3 in γc Cytokine Signaling | 0.418 | 0.029 | NaN |
| SPINK1 General Cancer Pathway | 0.418 | 0.029 | NaN |
| Role of JAK family kinases in IL-6-type Cytokine Signaling | 0.416 | 0.04 | NaN |
| Glutathione Redox Reactions I | 0.416 | 0.04 | NaN |
| Tryptophan Degradation X (Mammalian, via Tryptamine) | 0.416 | 0.04 | NaN |
| Role of JAK1, JAK2 and TYK2 in Interferon Signaling | 0.403 | 0.0385 | NaN |
| D-myo-inositol (1,4,5)-Trisphosphate Biosynthesis | 0.403 | 0.0385 | NaN |
| IL-10 Signaling | 0.396 | 0.0278 | NaN |
| Role of NANOG in Mammalian Embryonic Stem Cell Pluripotency | 0.393 | 0.025 | NaN |

### Supplemental Table 1, continued

| Ingenuity Canonical Pathways | -log(p-value) | Ratio | z-score |
|---|---|---|---|
| Dopamine Degradation | 0.368 | 0.0345 | NaN |
| Role of p14/p19ARF in Tumor Suppression | 0.368 | 0.0345 | NaN |
| B Cell Receptor Signaling | 0.367 | 0.0207 | -1.897 |
| G-Protein Coupled Receptor Signaling | 0.362 | 0.0217 | NaN |
| Dopamine Receptor Signaling | 0.361 | 0.026 | NaN |
| HER-2 Signaling in Breast Cancer | 0.357 | 0.022 | -2.236 |
| VDR/RXR Activation | 0.355 | 0.0256 | NaN |
| NF-κB Activation by Viruses | 0.355 | 0.0256 | NaN |
| Toll-like Receptor Signaling | 0.355 | 0.0256 | NaN |
| Thyroid Cancer Signaling | 0.348 | 0.0253 | NaN |
| GABA Receptor Signaling | 0.337 | 0.0229 | NaN |
| BAG2 Signaling Pathway | 0.319 | 0.0238 | NaN |
| Role of JAK2 in Hormone-like Cytokine Signaling | 0.317 | 0.0294 | NaN |
| Pyrimidine Ribonucleotides Interconversion | 0.317 | 0.0294 | NaN |
| Sirtuin Signaling Pathway | 0.311 | 0.0205 | 2 |
| DNA Methylation and Transcriptional Repression Signaling | 0.308 | 0.0286 | NaN |
| Pyrimidine Ribonucleotides De Novo Biosynthesis | 0.299 | 0.0278 | NaN |
| TWEAK Signaling | 0.292 | 0.027 | NaN |
| Superpathway of Methionine Degradation | 0.292 | 0.027 | NaN |
| Hereditary Breast Cancer Signaling | 0.29 | 0.0211 | NaN |
| Regulation of the Epithelial-Mesenchymal Transition Pathway | 0.288 | 0.0205 | NaN |
| Ceramide Signaling | 0.287 | 0.0222 | NaN |
| RANK Signaling in Osteoclasts | 0.281 | 0.022 | NaN |
| Antigen Presentation Pathway | 0.276 | 0.0256 | NaN |
| Factors Promoting Cardiogenesis in Vertebrates | 0.256 | 0.0199 | NaN |
| April Mediated Signaling | 0.255 | 0.0238 | NaN |
| B Cell Activating Factor Signaling | 0.248 | 0.0233 | NaN |
| BER (Base Excision Repair) Pathway | 0.242 | 0.0227 | NaN |
| iNOS Signaling | 0.224 | 0.0213 | NaN |
| nNOS Signaling in Skeletal Muscle Cells | 0.218 | 0.0208 | NaN |
