## Supplementary figures and images for "The aging lung mucosa: A proteomics study"

### Supplemental Figure 1

#### Supplemental Figure 1

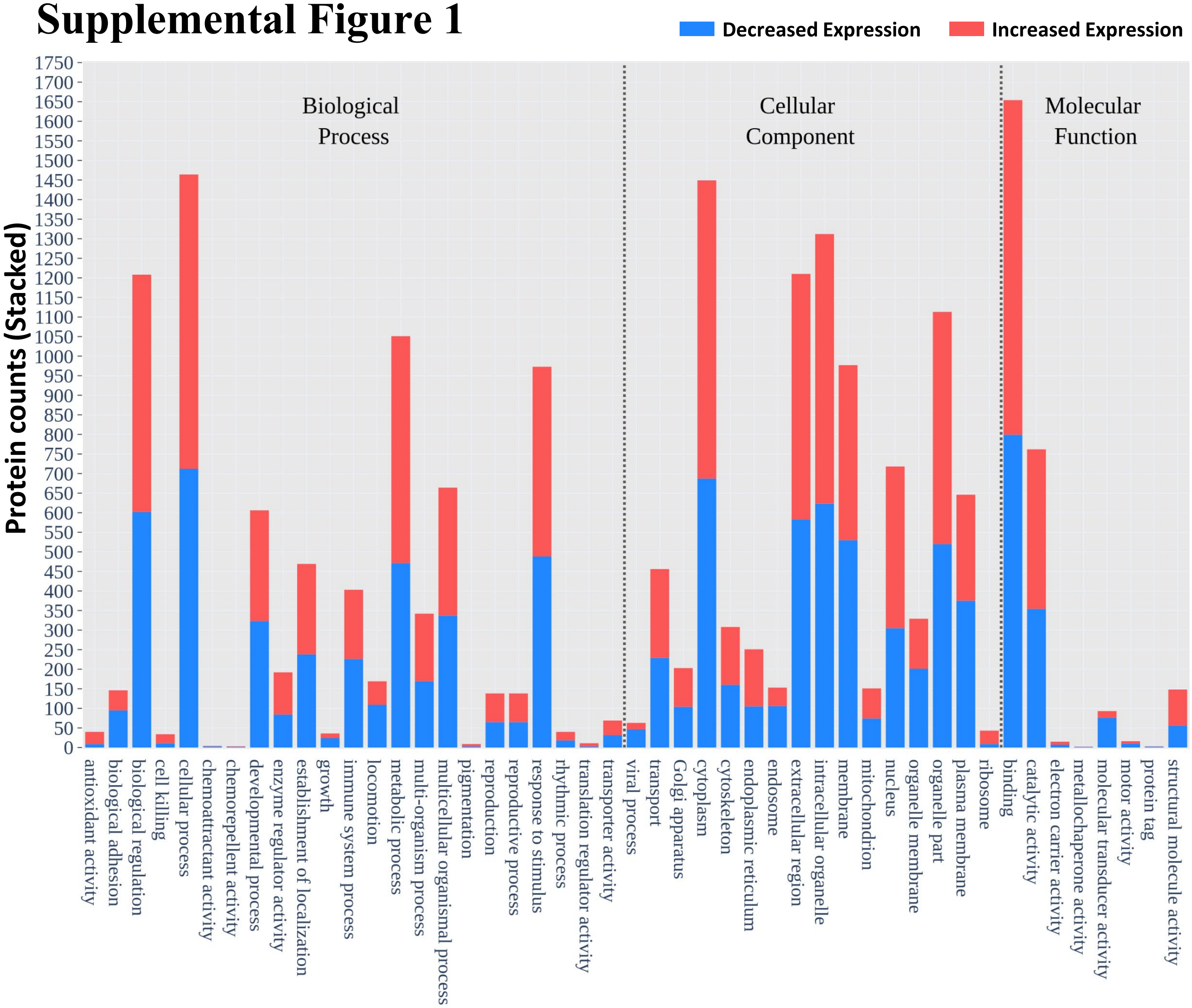
